# Transit amplifying cells balance growth and differentiation in above-ground meristems

**DOI:** 10.1101/2024.05.04.592499

**Authors:** Jessica Joossens, Denia Herwegh, Reinout Laureyns, Julie Pevernagie, Tom Van Hautegem, Lotte Pollaris, Samik Bhattacharya, Christian Korfhage, Thomas Depuydt, Kirin Demuynck, Klaas Vandepoele, Yvan Saeys, Clinton Whipple, Josh Strable, Hilde Nelissen

## Abstract

In both animals and plants, stem cell niches balance between cell-renewal and the generation of progeny cells that differentiate into specialized tissues. The multipotent and highly proliferative transit amplifying cells (TACs) integrate signals from stem cells and their differentiating progeny cells. Here we used spatial transcriptomics mapped to individual cells to illustrate the localization of TACs in maize meristems based on transcriptional gradients. Through genetic interactions and fluctuations in the transcriptional gradients we show that the multiplicative cell divisions are independently controlled from the TAC cell divisions. The dynamic nature of transcriptional variation in response to cell state or environment, together with the potential to improve yield by their modulation highlights the importance of finetuned modulation of networks rather than constitutive perturbations for crop improvement.

---

In multicellular organisms, stem cells are undifferentiated cells capable of self-renewal and differentiation into specialized cells. Plant and animal stem cells share commonalities because they are organized in specialized microenvironments or stem cell niches, which are part of meristems in plants. While animal stem cells are mostly considered dormant, cell divisions ramp up in cells at the transition between stem cells and differentiated cells, called transit amplifying cells (TAC), that account for the bulk of tissue production. TACs are not only a transitory phase from stem cells to post-mitotic cells, but they also actively control the number of daughter cells that transition towards tissue specific functions^1^.

In the plant shoot apical meristem (SAM), the organized structure that harbors the plant stem cell niche for shoot development and orchestrates continuous organ formation, there is currently no clear definition of cells equivalent to the animal TACs. In the maize SAM, mRNA *in situ* hybridizations (ISH) showed that *KNOTTED1* (*KN1*) expression marked meristematic cells but was absent from incipient primordia^2^, suggesting a rather sudden transition from meristematic cells to lateral organ growth and tissue differentiation. This lack of overlap between the expression domains of meristematic and differentiation markers, such as *KN1* and *YABBY* genes, respectively, is likely an incomplete view in light of a continuous cell differentiation state that was observed in single-cell (sc) RNA sequencing data from the maize SAM^3^. *PLASTOCHRON1* (*PLA1*) expression is at the boundary between meristematic and differentiating cells^4^, but the specific cells expressing *PLA1*, as well as the mechanistic insight into the role of *PLA1*, remains elusive. Here we study whether the cells expressing *PLA1* represent the plant counterpart of the TACs in the maize SAM.

## *PLASTOCHRON1* transcript accumulation overlaps with meristematic and differentiating cells

Cellular co-expression with *PLA1* was examined using single molecule fluorescence ISH (smFISH). Using a weighted gene co-expression network analysis of laser microdissection followed by RNA sequencing^5^, *PLA1* was positioned together with *KN1*, as well as with *GIF1* and the *YABBY* transcription factor encoding gene *YABBY9* (*YAB9*), which was confirmed by the co-expression tools CORNET^6^ and ATTED-II^7^ (**Supplemental figure 1**; **Supplemental table T1 and T2**). Based on these co-expression data, *PLA1, KN1, GIF1, YAB9* and its close homolog *YAB14*, together with specific markers of the SAM, including marker genes for the tip *ARGONAUTE18a* (*AGO18a*) and *DYNAMIN-RELATED PROTEIN4a* (*DRP4a)*, vasculature *LIKE AUXIN RESISTANT2* (*LAX2*) and *PIN*-*FORMED1* (*PIN1*), and boundary *CUPSHAPED COTELYDON2* (*CUC2*) (**Supplemental table T3**) were used as probes in the smFISH experiment of the maize shoot apex. The data can be visualized at https://www.psb.ugent.be/shiny/space_tool/ (**Username:** reviewer; **Password:** 6N89QKTL).

Simultaneous visualization of *PLA1, GIF1, KN1, YAB14* and *YAB9* expression confirmed co-expression with *PLA1*, albeit partially (**Figure 1A**). Cell segmentation (**Figure 1B**) and transcript read mapping to individual cells showed that the meristematic marker *KN1* and the differentiation markers *YAB14* and *YAB9* were not co-expressed at the boundary between the meristem and the leaf primordia (**Figure 1C-D**). Only few cells co-expressed *KN1* with *YAB14* and *YAB9* in the rib zone and in primordia cells (0.65% and 0.52%, respectively; **Supplemental table T4**). The lack of overlap between the *KN1* and *YAB* expression domains was suggested in single gene ISHs^2,8,9^, while the overlap between *KN1* and *YAB14* and *YAB9* was also minimal in maize SAM scRNA sequencing data (**Supplemental figure S2A-D**;^3^). *PLA1* was co-expressed with both meristematic and differentiating markers because 59% of the cells expressing *PLA1* also expressed *KN1* (**Figure 1E**), while cells, located at the base of young leaf primordia, co-expressed *PLA1* with *YAB14* (4%) (**Figure 1F**) and with *YAB9* (2%) (**Supplemental figure S3A**). *GIF1* was expressed in almost every cell where *PLA1* was expressed (92% overlap, **Supplemental figure S3B**) and up to 19% and 5.5% of cells that expressed *GIF1* also expressed *YAB14* (**Figure 1G**) or *YAB9* (**Supplemental figure S3C**), respectively. In the rib zone, *GIF1* is co-expressed with *KN1*, except in vascular cells where *GIF1* expression is absent (**Figure 1H**). The co-expression was also seen in the scRNA sequencing data from the maize shoot apex (**Supplemental figure S2A, C-F**;^3^). In summary, *PLA1* is locally co-expressed with meristematic and differentiating genes, whereas *GIF1* is more broadly co-expressed with meristematic and differentiation markers, as well as with *PLA1*.

**Figure 1:**
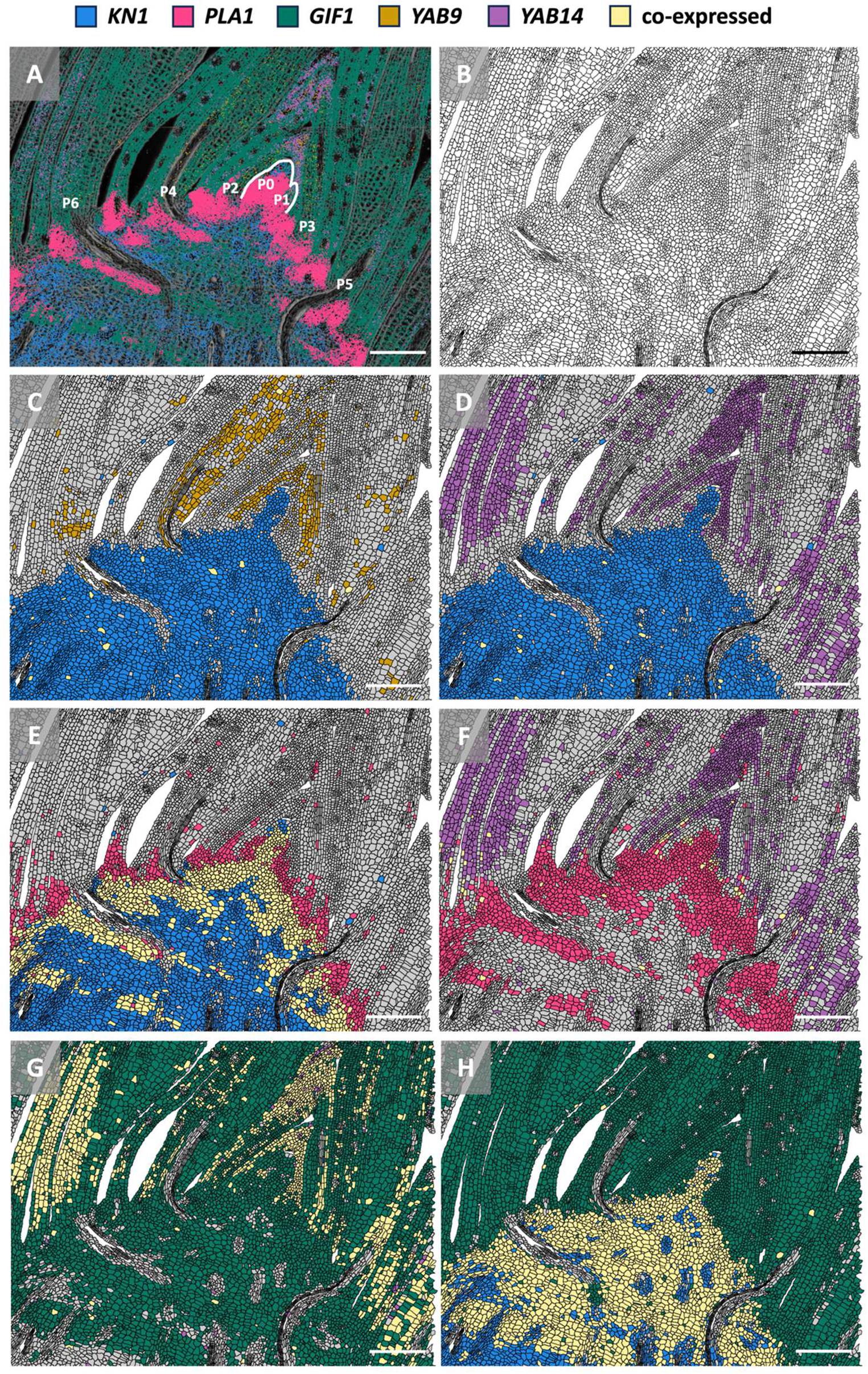
Pairwise comparison of *KN1, PLA1, GIF1, YAB9* and *YAB14* in the maize shoot apex. (**A**) Visualization of *KN1*, *PLA1*, *GIF1*, *YAB9* and *YAB14* expression. The white line indicates the shoot apical meristem including the incipient primordia 0 (P0) and P1. (**B**) Cell segmentation of the shoot apex. (**C-H**) Co-expression between (**C**) *KN1* and *YAB9*, (**D**) *KN1* and *YAB14*, (**E**) *KN1* and *PLA1*, (**F**) *PLA1* and *YAB14*, (**G**) *GIF1* and *YAB14* and (**H**) *KN1* and *GIF1*. Scale bars = 200 µm. Molecular cartography was used to generate the spatial maps.

## Spatially resolved cellular co-expression positions *PLASTOCHRON1* in a transcriptional gradient between undifferentiated and differentiating cells

Clustering the single cells according to the transcriptional landscape, followed by a dimensionality reduction (**Figure 2A-B**) facilitated cell type specific co-expression. At the tip of the SAM (cluster 8), *KN1* and *GIF1* are co-expressed whereas co-expression of *PLA1* with *KN1* and *GIF1* is restricted to cells that subtend the meristem tip (**Supplemental figure S4A-C**) and the boundary region between meristematic and differentiating cells (cluster 3; **Supplemental figure S4D-E**). Cells extending into the base of young leaf primordia (cluster 3) expressed *YAB14* and *YAB9* together with *PLA1* (**Supplemental figure S4F-G**), while *KN1* expression is absent. Cells at the base of leaf primordia (cluster 7) co-expressed *PLA1* and *GIF1*, but *PLA1* is no longer co-expressed with *KN1* (**Supplemental figure S4H-I**). More distal from the leaf base (cluster 6), differentiating cells are marked by high expression of *GIF1*, *YAB14* and *YAB9* and co-expression between *GIF1* and *YAB14* (**Supplemental figure S4J-K**). These data show that coordinated expression between *KN1*, *PLA1 GIF1*, *YAB14* and *YAB9* is cell type dependent.

**Figure 2:**
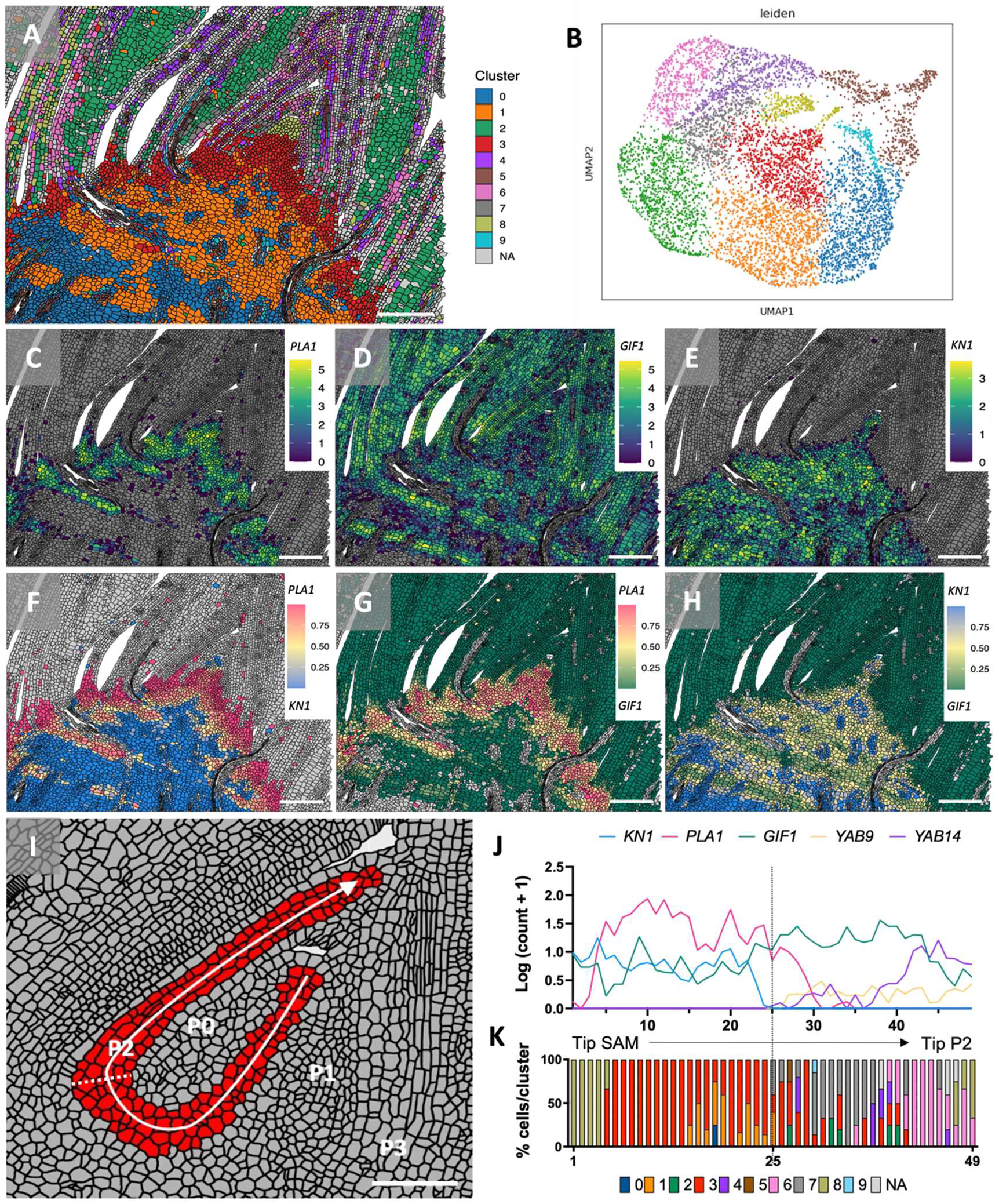
Quantitative expression analysis of *KN1, PLA1, GIF1, YAB9* and *YAB14* in the maize shoot apex. (**A**) Dimensionality reduction and cell classification. Scale bar = 200 µm. (**B**) UMAP plot. (**C-E**) Expression gradients of (**C**) *PLA1*, (**D**) *GIF1* and (**E**) *KN1* in the shoot apex using the log-transformed counts per cell. (**F-H**) Pairwise quantitative comparison between (**F**) *KN1* and *PLA1*, (**G**) *GIF1* and *PLA1* and (**H**) *KN1* and *GIF1*. Scale bars = 200 µm. (**I**) Selected cells (red) for the predefined spatial trajectory going from the tip of the SAM to the tip of P2 (white arrow). Scale bar = 100 µm. (**J**) Transcriptional gradients of *KN1, PLA1, GIF1, YAB9* and *YAB14* along the spatial trajectory. Transcript counts per cell were log-transformed and the average of 3-5 horizontally aligned cells are shown along the predefined spatial trajectory. (**K**) Percentage of cells per cluster along the spatial trajectory. Dotted line indicates the clear delineation between *KN1*, the meristematic marker, and *YAB9* and *YAB14*, the differentiation markers. Molecular cartography was used to generate the spatial maps.

To understand quantitative dynamics in transcript accumulation, expression gradients were visualized. *PLA1* transcript accumulation was highest in regions subtending leaf primordia and gradually lowered towards differentiating cells and the SAM tip^4^ (**Figure 2C**). *PLA1* expression corresponded to regions of high cell division activity, as indicated by the co-expression with *HISTONE H4*^4^ (**Supplemental figure S5**) and with regions where the cell cycle markers *HISTONE H3* and *CYCLIN1* were expressed^3^. *GIF1* expression was relatively low in the SAM and meristematic cells but increased towards older intercalary meristems and differentiating cells (**Figure 2D**), corresponding with a role in stem cells and differentiating cells^10,11^. *KN1* expression gradually decreased towards more developed vasculature and differentiating leaf cells (**Figure 2E**). An expression gradient from meristematic cells to early differentiating cells was visualized when combining the quantitative expression of *KN1* and *PLA1* (**Figure 2F**). Pairwise comparison of *PLA1* with *GIF1* (**Figure 2G**) highlights an almost complementary transcript accumulation, with a lower *GIF1* expression in the boundary cells at the base of the developing primordia, where *PLA1* expression was maximal. Both *PLA1* and *GIF1* were highly co-expressed in meristematic cells, but *PLA1* was predominantly expressed in early intercalary meristems, whereas *GIF1* was predominantly expressed in older intercalary meristems. *GIF1* expression was higher than that of *KN1* in older intercalary meristem cells (**Figure 2H**), whereas *KN1* expression was higher in developing vasculature and in the rib zone. So, even in cells that co-express *KN1*, *PLA1* and *GIF1*, quantitative differences determined a spatial and temporal transcriptional gradient.

Gene counts of *KN1*, *PLA1*, *GIF1*, *YAB9* and *YAB14* were quantified per cell and averaged for 3-5 aligned cells across a predefined spatial trajectory from the tip of the SAM to the P2 (**Figure 2I-K**), confirming the clear delineation between *KN1* and *YAB14 and YAB9*. Around the P0 and P1 primordia, where a decrease of *KN1* expression was necessary for leaf initiation^2^, we see increased expression counts of *GIF1*. *PLA1* expression gradually increased from the tip of the SAM towards the rib zone. In cells closer towards leaf primordia, marked by loss of *KN1* expression and initiation of *YAB14* expression, *PLA1* rapidly declined, accompanied by a steady increase in *GIF1* expression. *PLA1* expression leveled out several cell files after *KN1* expression disappeared, while *PLA1* expression was maximal around P0 and at the leaf base, two boundary regions between meristematic cells and differentiating leaf cells. These transcriptional gradients were also observed in spatial trajectories going from the tip of the SAM to P0, P1 and P3 (**Supplemental figure S6**). In a developmental trajectory tracing cell differentiation in the SAM using scRNA sequencing, similar dynamic expression patterns were seen for *KN1*, *PLA1*, *GIF1*, *YAB9* and *YAB14*, where the genes clustered separately over pseudotime^3^. Together with the PLA1 stimulatory role of cell division activity^12^, the *PLA1* expression in both meristematic and differentiating cells, that co-localizes with active cell divisions suggests a role for *PLA1* expressing cells to be the plant counterpart of TACs.

## PLASTOCHRON1 is involved in the maintenance of transit amplifying cells

To investigate if PLA1 is involved in TACs, we examined SAM morphology upon *PLA1* perturbation. The area of the SAM was increased in the *pla1-2* mutant compared with wild type (+44%, P ≤ 0.05; **Figure 3A-C**), mainly due to an increase in SAM height (+32%, P ≤ 0.05; **Figure 3D-E**). Therefore, PLA1 not only controls the size of differentiated organs^12^, but also meristem size. The number of cells expressing *DRP4a*, a marker for the tip of the SAM^13^, was decreased in the *pla1-2* mutant (−58%, P ≤ 0.01; **Figure 3F-I**), indicating more cells are drained away from the stem cell niche into differentiation in the *pla1-2* mutant. The *pla1-2* mutant had fewer *PLA1* expressing cells than wild type (−41%, P = 0.11; **Figure 3J-L**), albeit not significant, but the normalized *PLA1* transcript counts were significantly decreased in the *pla1-2* mutant (−57%, P ≤ 0.05; **Figure 3M**), suggesting that PLA1 regulates its own expression level. To test if PLA1 functioned in a dose-dependent manner, the *pla1-2* mutant was crossed with the *GA2ox:PLA1* line (**Figure 3N**). The complementation analysis revealed a dosage dependent effect of PLA1 on leaf length, elongation rate and duration, with *pla1-2;GA2ox:PLA1* plants performing better than wild type but not the *GA2ox:PLA1* plants (**Figure 3O-Q; Supplemental table T5**).

**Figure 3:**
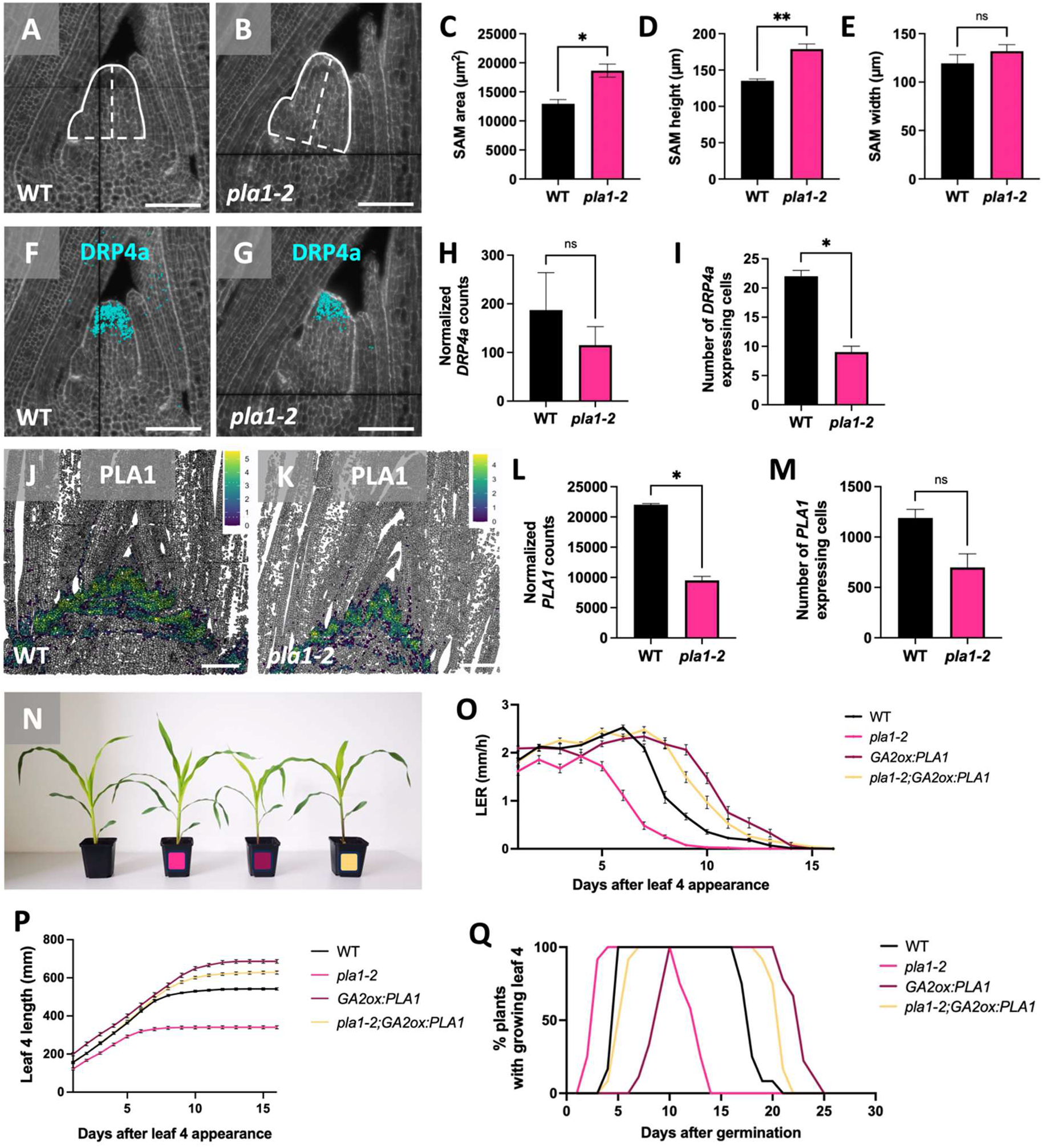
PLA1 is involved in the maintenance of transit amplifying cells. Shoot apical meristem (SAM) morphology in (**A**) wild type and (**B**) the *pla1-2* mutant. Phenotypes for (**C**) SAM area (white line), (**D**) height (vertical segmented line) and (**E**) width (horizontal segmented line). Significance was calculated using a Welch’s t-test (n = 3). *DRP4a* expression in the SAM of wild type and the *pla1-2* mutant. (**H**) Normalized *DRP4a* counts in the SAM and (**I**) the number of cells expressing *DRP4a*. Significance was calculated using a Welch’s t-test (n = 2). Log-transformed *PLA1* expression in (**J**) wild type and (**K**) the *pla1-2* mutant. (**L**) Normalized *PLA1* counts in the shoot apex and (**M**) the number of cells expressing *PLA1*. Significance was calculated using a Welch’s t-test (n = 2). (**N-Q**) Double mutant analysis of a segregating *pla1-2;GA2ox:PLA1* population. (**N**) From left to right: segregating wild type, *pla1-2* (pink dot), *GA2ox:PLA1* (dark red dot) and *pla1-2;GA2ox:PLA1* (yellow dot) plants, all germinated on the same day. Phenotypes for (**O**) leaf elongation rate (LER), (**P**) leaf 4 length and (**Q**) the percentage of plants over time with a growing leaf four. The significance of the LER was calculated using a mixed model with Bonferroni correction (n ≥ 9). Leaf 4 length was calculated using a one-way analysis of variance with Bonferroni correction (n ≥ 9). Statistical analysis for leaf 4 length can be found in **Supplemental table T5**. * = P ≤ 0.05 and ** = P ≤ 0.01. ns = non-significant. Molecular cartography was used to generate the spatial maps.

Because PLA1 affects meristem size, we evaluated whether KN1, a putative regulator of *PLA1* and *GIF1*^14^, affected the genes expressed in the TACs and *vice versa*. *PLA1* and *GIF1* expression was not obviously altered in the dominant, gain-of-function *KN1-O* and *KN1-DL* mutants^15^ nor in the loss-of-function mutant *kn1*-*e1*^16^ (**Supplemental figure S7A-N**). Few target genes of KN1 are perturbed in *kn1* mutants^13^, which could explain the lack of changes in *PLA1* and *GIF1* expression. Spatial maps of the shoot apex for lines carrying perturbations in *PLA1* revealed no changes in *KN1* expression counts in the *pla1-2* mutant (**Supplemental figure S8A-B**), but between *GA2ox:PLA1* and control wild type plants, a 2.7-fold increase in expression counts for *KN1* was detected (**Supplemental figure S8C-E**). A similar threefold increase was also observed in a transcriptome analysis of the basal half cm of leaf 4 from *GA2ox:PLA1* transgenic plants^12^ (**Supplemental figure S8F**). Conversely, no effect on *KN1* expression was observed in the *pla1-2* or *gif1-2* mutants compared with the respective wild type segregants using ISH (**Supplemental figure S9A-D**).

## PLA1 and GIF1 independently coordinate TAC and multiplicative cell divisions

Because almost all *PLA1* expressing cells express *GIF1*, GIF1 could be part of TAC regulation, but the broader expression domain of *GIF1* could also suggest a more general role for GIF1. The *gif1* mutants displayed both meristem determinacy and leaf growth phenotypes^10^ and no growth effect was observed when overexpressing *GIF1* using the constitutive ubiquitin promoter^11,17^. Because *GIF1* is highly expressed^13^, a stronger *Brachipodium distachyon EF1α* promoter^18^ was used to achieve higher levels of overexpression (**Figure 4A, Supplemental figure 10A**). Only *EF1α:GIF1-07* (*GIF1-07*) with the highest *GIF1* overexpression level displayed a significant increase in final leaf length (+8.6%, P ≤ 0.001) (**Supplemental figure S10B**), similar to what was seen in Arabidopsis^19^. This growth phenotype was due to a significant increase in the duration of leaf elongation (**Figure 4B**, **Supplemental table T6**) and cell division activity (**Figure 4C**). The increased leaf growth persevered until plants reached maturity in the greenhouse (**Supplemental figure S10C-D**) and in the field (**Supplemental figure S10E**). Strong overexpression of *GIF1* in the *EF1α:GIF1-07* line also positively affected plant biomass and seed yield in field trials (**Supplemental figure S10E-F; Supplemental table T7**). Stem width and leaf biomass were increased over all field trials, leading to an overall increase in plant fresh weight of the *GIF1-07* line in B104 inbred (+28%, P ≤ 0.01) and *GIF1-07/*B104 x CML91 hybrid (+25%, P = 0.063) background. Seed yield was increased in the field, with average cob weight increased in two field trials (P ≤ 0.05), due to increased cob length (+10%, P ≤ 0.001) and 100 kernel weight (+23%, P ≤ 0.001). In the dry season of 2022, cob weight was not significantly different, but *GIF1-07 x CML91* hybrids had a 17% increase in 100 kernel weight (P ≤ 0.001) compared with the non-transgenic hybrids. The cellular mechanism and the positive effect on biomass and seed yield of *GIF1* overexpression resembled what was seen for ectopic expression of *PLA1*^20^, providing indication that *PLA1* and *GIF1* could function together.

**Figure 4:**
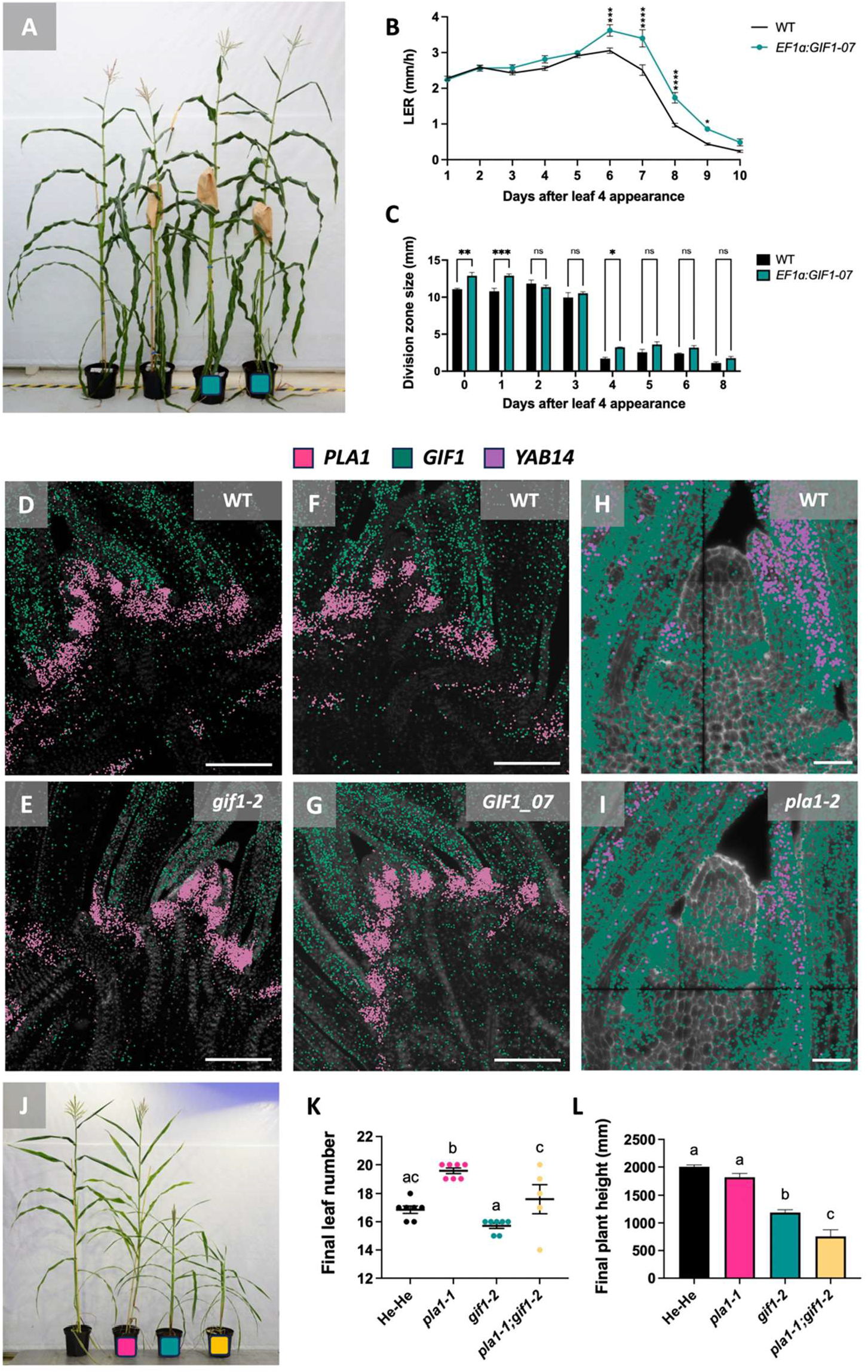
PLA1 and GIF1 independently coordinate TAC and multiplicative cell divisions. (**A**) Phenotypes of the *EF1α:GIF1-07* transgenic line. From left to right: two segregating wild type plants and two *EF1α:GIF1-07* siblings (marked by green dot). (**B**) The leaf elongation rate (LER) and (**C**) the division zone size (DZS) of leaf 4 in segregating wild type and *EF1α:GIF1-07* plants. Significant differences for LER were calculated by a mixed analysis of variance with Bonferroni correction (n ≥ 5). Significant differences for DZS were calculated by a two-way analysis of variance with Bonferroni correction (n = 3). (**D-E**) Expression of *PLA1* and *GIF1* in the shoot apex of (**D**) wild type and (**E**) the *gif1-2* mutant using *in situ* sequencing. Scale bars = 200 µm. (**F-G**) Expression of *PLA1* and *GIF1* in the shoot apex of (**F**) wild type and (**G**) the *EF1α:GIF1-07* overexpression line using *in situ* sequencing. Scale bars = 200 µm. (**H-I**) Expression of *GIF1* and *YAB14* in (**H**) wild type and (**I**) the *pla1-2* mutant using molecular cartography. Scale bars = 50 µm. (**J**) From left to right: The *pla1-1*/+;*gif1-2*/+ control, *pla1-1* (marked by pink dot), *gif1-2* (marked by green dot) and *pla1-1;gif1-2* mutant (marked by yellow dot). (**K-L**) Phenotypes of the *pla1-1;gif1-2* plants, including (**K**) final leaf number and (**L**) final plant height. Significance was calculated by an analysis of variance with Bonferroni correction (n ≥ 5).

To address whether *PLA1* and *GIF1* influence each other’s transcription, spatial maps were generated of overexpression and mutant lines. *PLA1* was expressed more widely in the SAM of *gif1-2* mutants (**Figure 4D-E**) and was diminished in the intercalary meristem of *GIF1-07* overexpression lines (**Figure 4F-G**; **Supplemental figure S11A-D**). In the *pla1-2* mutant, two bands of *GIF1* expression were present, marking new incipient primordia (P0), of which only one was present in the control at the same developmental stage (**Figure 4H-I; Supplemental figure S11E-F**), confirming the higher rate of leaf initiation that was seen in the *pla1-2* mutant^4^. In addition, *GIF1* expression showed a 41% decrease in expression counts over the entire shoot apex of *GA2ox:PLA1* (**Supplemental figure S12A**). Conversely, a transcriptome analysis of the basal half cm of leaf 4, enriched for dividing cells from *GA2ox:PLA1*^12^ revealed *GIF1* upregulation, which was confirmed using RT-QPCR (**Supplemental figure S12B**). The opposing effects of *PLA1* perturbation on *GIF1* expression in different organs was in line with their spatiotemporal and tissue-specific expression. These data showed that *PLA1* and *GIF1* expression was altered in the respective mutants, but were not conclusive on whether this was due to the fact that PLA1 and GIF1 influenced each other’s expression or that the changes in morphology were reflected in altered expression.

To investigate the genetic interaction between *PLA1* and *GIF1*, we performed a double mutant analysis with the *pla1-1* mutant^20^ and the *gif1-2* mutant^10^ (**Figure 4J**). *pla1-1;gif1-2* double mutant plants produced more leaves compared with wild type and single *gif1* mutant plants but less leaves than in the *pla1-1* mutant (**Figure 4K**). *pla1-1;gif1-2* double mutants exhibited a decreased plant growth of 62.2% compared with the wild type siblings (P ≤ 0.001), which was more than the decrease in either single mutant (9.4% for *pla1-1* and 40.7% for *gif1-2*; **Figure 4L**). Similarly, the leaf lengths of the *pla1-1;gif1-2* double mutants were significantly decreased compared with those of wildtype and *pla1-1* plants for all leaves, while this decrease was only significant for leaves 9 and 10 when compared with the *gif1-2* mutants (**Supplemental figure S13, Supplemental table T8**). About half of the double mutant plants did not develop functional reproductive organs (**Supplemental figure S14**). The single *gif1-2* mutant was male and female sterile^10^, while the *pla1-1* mutant developed reproductive organs, but with reduced seed set. Together, the phenotypes of the *pla1-1;gif1-2* double mutants suggest an additive interaction between *PLA1* and *GIF1*, meaning that the two pathways to stimulate cell division act independently and that the change in expression profiles in the respective mutants is mainly due to the anatomical changes. These data show that the cell division processes in the TACs coordinated by PLA1 are different from the multiplicative cell divisions controlled by GIF1.

## Transit amplifying cells are conserved in the vascular procambium and inflorescence meristems

Although the majority of *PLA1* expression was at the boundary region, more subtle *PLA1* expression was seen in young leaf primordia (**Figure 2C and 2F**) as well as along the gradient of a growing leaf four^12^ (**Figure 5A, white square**), suggesting that the *KN1-PLA1-GIF1* module might not be limited to the SAM. In young leaf primordia, *PLA1* expression was present in the ground meristem and the procambium cells of the developing 2^°^ and 3^°^ veins of P3 and P4 (**Figure 5A**, **blue square**), which is in accordance with a laser capture microdissection followed by RNA sequencing experiment^21^. *KN1* is co-expressed with *PLA1* and *GIF1* in the provascular cells of the rib zone which are marked by *LAX2* expression (**Supplemental figure 15**). LAX2, an auxin transporter that was identified as a marker of the ground meristem^21^, was co-expressed with *PLA1* (**Figure 5B**), more specifically at the 3^°^ veins of P3 and P4 (**Figure 5C**) and in the provascular cells of the rib zone. Moreover, high *PLA1* expression was seen at the abaxial side of the developing midvein in young leaf primordia (**Figure 5A, white square**), comprising the protoderm and the sub-protodermal ground meristem. In the *pla1-2* mutant, Kranz anatomy and sclerenchyma cells were differentially organized compared with wild type (**Figure 5D-E**) as there were less and smaller sclerenchyma cells at the abaxial side of the midvein (P ≤ 0.0001; **Figure 5F**). These data show that *PLA1* expression is activated whenever new cell divisions are needed, during organ formation from the SAM but also during new tissue generation, indicating a more general role for the TACs. *KN1*, *PLA1*, *GIF1*, *YAB9* and *YAB14* expression was also present in the developing ear (**Figure 5G-H**), with similar transcriptional gradients in ear spikelet meristems as in the SAM (**Supplemental figure S16**). The highest *KN1* expression was at the tip of the spikelet meristem, while *YAB9* and *YAB14* were expressed in the lateral bract and glume and with no overlap between *KN1* and *YAB* expression. *GIF1* was expressed in both the meristematic region as well as in the palea, lower floret meristem and glume, while *PLA1* was expressed at the boundary between developing spikelets, showing that TACs are present in various above-ground meristems to coordinate growth and differentiation.

**Figure 5:**
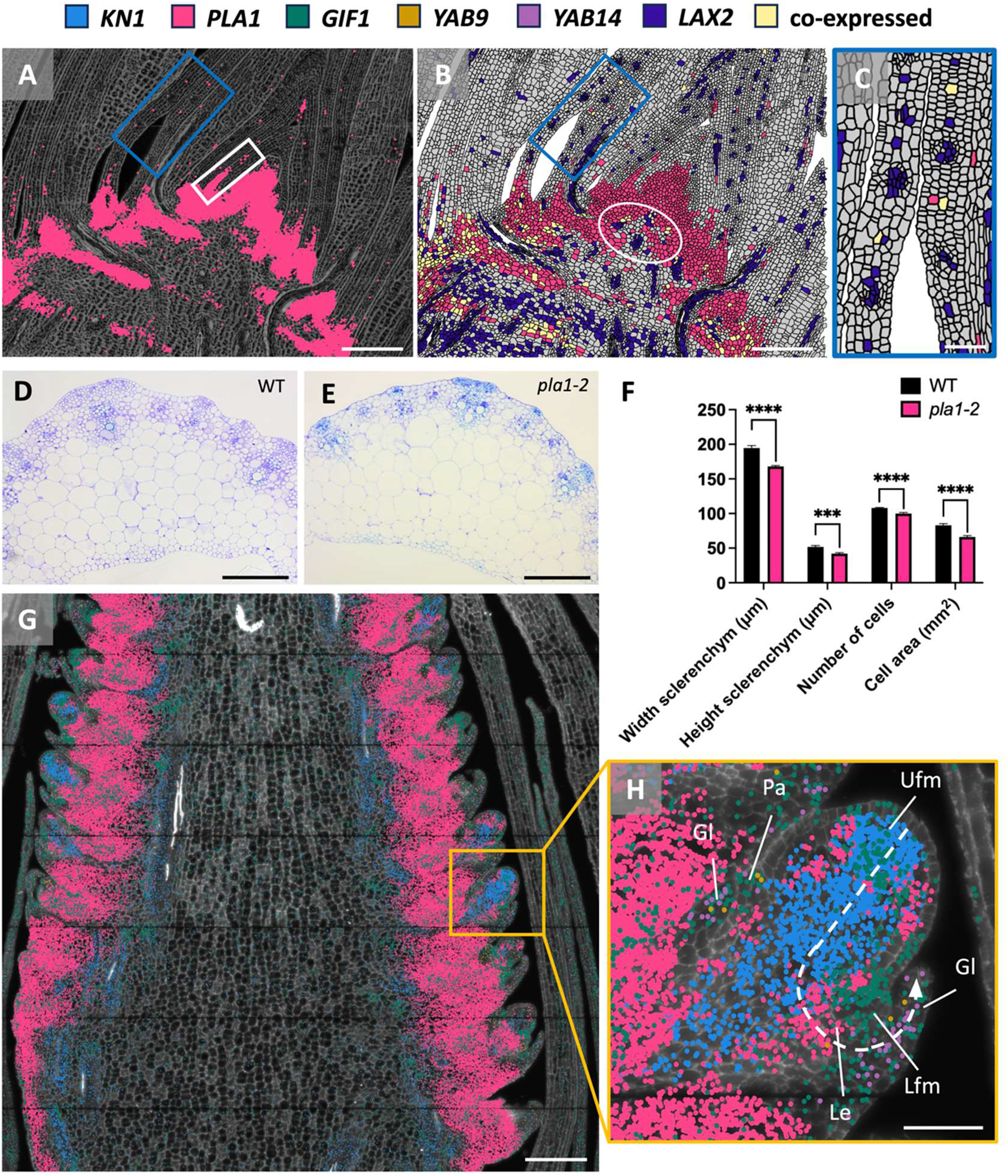
Transit amplifying cells are conserved in the vascular procambium and inflorescence meristems. (**A**-**C**) Visualization of *PLA1* and *LAX2* in the maize shoot apex. (**A**) *PLA1* is expressed in 2^e^ and 3^e^ veins of P3 and P4 (blue square) and in the cell files abaxial from the midvein (white square). Scale bar = 200 µm. (**B**) Co-expression of *PLA1* and *LAX2* in the 2^e^ and 3^e^ veins of P4. Scale bar = 200 µm. (**C**) Zoom in of B on P4 margins: *PLA1* and *LAX2* are co-expressed in the procambium cells of the 3^e^ veins (yellow-colored cells). Scale bar = 50 µm. (**D**-**E**) Transversal section of a growing leaf 4 showing differences in kranz anatomy and sclerenchyma cells between (**D**) wild type and (**E**) the *pla1-2* mutant. Scale bar = 200 µm. (**F**) Phenotypes in sclerenchyma cell size and number of cells in the *pla1-2* mutant compared with wild type. Significance for each phenotype was calculated by a two-tailed t test (n ≥ 23). **** = P ≤ 0.0001 and *** = P ≤ 0.001. (**G**) Visualization of *KN1*, *PLA1*, *GIF1*, *YAB9* and *YAB14* in a young maize ear (∼10 mm). The yellow square indicates the spikelet used for the predefined spatial trajectory (**Supplemental figure S16**). Scale bar = 200 µm. (**H**) Zoom in of G: visualization of the predefined spatial trajectory (white dashed arrow) in the ear spikelet, going from the tip of the upper floral meristem (UFM) to the tip of the glume (Gl). Lfm: lower floral meristem, Pa: palea, Le: lemma. Scale bar = 50 µm. Molecular cartography was used to generate the spatial maps.

## Conclusions

Although TACs have been reported in plants^3,22–24^, transcriptional dynamics with cellular resolution illustrated that in the maize SAM, TACs can rather be distinguished based on their molecular profiles than on their anatomical properties. Because PLA1 belongs to the plant-specific CYP78A family of cytochrome P450 monooxygenases, plant TACs might have evolved to cope with the continuous formation of new organs from plant meristems in a highly coordinated spatiotemporal manner.

## Acknowledgements

The authors thank Laurens Pauwels, Stijn Aesaert and Griet Coussens for the maize transformation, Stan Werbrouck for maintance of the SPACE app, Julie Merchie and Bernard Cannoot for the greenhouse phenotyping, and all field volunteers for the field measurements and the detasseling. The spatial transcriptomics approaches received funding from the VIB Tech Watch fund and support from Resolve Biosciences and Cartana. We also thank Jane Langdale and Chiara Perico for feedback on the manuscript.

## Material and methods

### Plant materials and background used in this study

*Zea mays* (maize) B104 background was used for generating the *GA2ox:PLA1* line (Sun et al., 2017), the *pla1-2* mutant (Laureyns et al., 2022) and the *EF1α:GIF1* lines in this study. The *pla1-1* mutator transposon line (mu1044329) is in a W22 background. Both *gif1-2* in a W22 and in B73 background were used for crosses with the *pla1-1* mutant. Heterozygous offspring was selfed for one or two generations to obtain a segregating population for double mutant analysis. For *in situ* sequencing (ISS) and in situ hybridization (ISH), the *GA2ox:PLA1*, *pla1-2*, *EF1α:GIF1-07* and *gif1-2* lines and their respective wild type controls were used. For molecular cartography and ISH, B104 wild type, the *pla1-2* mutant and the segregating wild type control were used. For ISH, the *KN1-DL* (Lisch et al., 1995), *KN1-0* (Ramirez et al., 2009) and the *kn1-e1* mutant in a permissive background (Kerstetter et al., 1997) were used.

### Maize transformation and genotyping

The maize GIF1 (Zm00001eb056300) was ligated into the vector pDONR221 using Gateway cloning (https://gatewayvectors.vib.be/) and combined with the *BdEF1*α promoter ^18^ in the vector pBbm24GW7. Immature embryos of the maize inbred line B104 were transformed by Agrobacterium tumefaciens cocultivation. We obtained independent events from transformation (*GIF-02*, *GIF-05* and *GIF-07*) with *GIF-07* showing the highest overexpression and phenotypic effects. Primary transgenic events in which the T-DNA was present in a single locus were backcrossed to B104 resulting in a 1:1 segregation. These segregating plants were used for the growth chamber and greenhouse evaluations in order to exclude maternal effects. For field trials, both transgenic and non-transgenic plants of the segregating generation were selfed (for two generations). The homozygous plants were crossed to CML91 to obtain transgenic and non-transgenic hybrids. The presence of the phosphinothricin acetyltransferase (PAT) protein was tested by an immunochromatographic assay (AgroStrip, Romer), leaf painting or PCR. Expression levels were monitored using RT-qPCR using 18S rRNA as housekeeping gene and the levels were determined using the 2 ^-ΔΔCT^ method. Primers used for genotyping and cloning are listed in **Supplemental table T9**.

### Growth conditions in growth chamber and greenhouse

For leaf growth monitoring on seedlings were grown under growth chamber conditions with controlled relative humidity (55%), temperature (24 °C day/18 °C night), and light intensity (170–200 μmol per m2 per second photosynthetic active radiation at plant level) provided by a combination of high-pressure sodium vapour (RNP-T/LR/400W/S/230/E40; Radium) and metal halide lamps with quartz burners (HRI-BT/400W/D230/E40; Radium) in a 16 h/8 h (day/night) cycle.

Plants for end point trait characterization were grown under controlled greenhouse conditions (26 °C/22 °C, 55% relative humidity, light intensity of 180 mmol per m2 per second photosynthetic active radiation, in a 16 h/8 h day/night cycle).

### Phenotyping in the growth chamber and greenhouse

Plants were measured daily from leaf emergence to growth arrest to determine the LER. Kinematic analysis was performed based on Nelissen et al., 2015. To determine the division zone size over time, leaf four was harvested daily before emergence from the sheath of leaf three until fully grown. The time point was determined by the day when leaf four was initially visualized from the whorl of leaf three. The size of the division zone was determined by the distance between the base and the most distally observed mitotic cell in DAPI-stained leaves along the proximal-distal axis with a fluorescence microscope (AxioImager, Zeiss). For every analysis, at least three technical replicates were taken.

### Field trial design and plant trait analysis

Field experiments were conducted from 2020 to 2022 at Wetteren, Belgium. In the first year, *GIF1-07* in B104 background and its non-transgenic control were grown. In 2021 & 2022, *GIF1-07 x CML91* and their non-transgenic controls were grown. They were grown in randomized block design with three replicates with planting density over 88,000 plants per hectare. Each replicate contained four rows and 40 plants per row. Vegetative trait analysis and biomass was measured on 20 representative plants per replicate. All cobs per replicate were photographed and measured for cob length using ImageJ. More detailed cob components (e.g. 100 kernel weight) were determined by harvesting 5 representative ears per replicate. Due to the late flowering of the B104 line, few cobs were pollinated which hampered cob analysis. The dry season of 2022 impacted growth and seed yields. Because the legislation in Belgium requires detasseling of transgenic plants in the field, both the transgenic and non-transgenic plants were detasseled and pollination originated from B104(xCML91) border plants. The necessity to detassel transgenic plants precluded any observation of anthesis, so that the anthesis silking interval could not be determined.

### Tissue collection, embedding and sectioning

Two days after leaf 4 appearance maize seedlings were harvested to collect shoot apex samples for *in situ* hybridizations and molecular cartography as described in Laureyns et al., 2022. Greenhouse grown B104 ears were harvested when they reached a length of approximately 10 mm (6 weeks after pollination). Samples were embedded in paraffin and sectioned in 7 µm thick sections using a microtome (Reichert-Jung 2040).

### mRNA *in situ* hybridization

Tissues were harvested and processed as described in Laureyns et al., 2022. ISH protocol was performed as described by Zöllner et al., 2021. Mounted sections were imaged using an Olympus BX51 light microscope. Primers used for probe generation are listed in **Supplemental table T9**. ISH were repeated at least twice.

### *In situ* sequencing

Section preparation, probe hybridization, ligation, amplification and visualization for ISS as described in Laureyns et al., 2022. At least 2 or 3 biological sections were available per genotype. Counts are normalized per section by dividing with all expression counts (normalized counts = counts per gene / all transcript counts).

### Section preparation for molecular cartography

In total, three biological repeats were obtained for longitudinal sections of the shoot apex for molecular cartography. Paraffin embedded sections were baked for 2 hours at 60 degrees for tissues to adhere to the slides. The sections were deparaffinized by 2x 30 minutes 100% histoclear steps. A gradient was applied to remove the histoclear and replace it with EtOH, 1 minute for each step. Finally, sections were brought in an aqueous solution by removing EtOH through a gradient, again 1 minute for each step. Rehydrated sections received a proteinase treatment 122 (Pronase: Sigma Aldrich) for 15 minutes which was stopped by placing sections in 0.2% (w/v) glycin dissolved in PBS. Sections were dehydrated through a gradient to 100% EtOH conditions. Slides were prepared for transportation to Resolve Biosciences by coating with SlowFade™ Gold Antifade mountant (Invitrogen). The genes were selected using ATTED and CORNET, co-expression with PLA1, correlation coefficient ≥ 0.7 and p-value ≤ 0,05 for CORNET and p-value of at least 0.01 for ATTED-II.

### Data analysis for molecular cartography

We benchmarked molecular cartography for use in plants using FFPE tissue sections of the maize shoot apex (Laureyns et al., 2022). The quality and reproducibility of the signals was high across three biological replicates (R² ≥ 0.96). Sensitivity and specificity were also high, with genes showing the expected expression pattern (**Supplemental table T3**) and low false positive rates (0.20%, 0.58%, 5.8%). Molecular cartography images were visualized in ImageJ using genexyz Polylux tool plugin from Resolve BioSciences. To quantify the expression on a cellular level, images obtained from the DAPI and calcofluor white stain were merged. Cells were segmented using CellPose (Stringer et al., 2021) with manual curation in QuPath (Bankhead et al. 2017) to train the model. In total, 11,795 cells were segmented. The segmented cells were saved as regions of interest (ROI), and gene expression counts of the 21 genes were allocated to the individual ROI’s. 54% of the cells have more than 20 counts and 326 cells (2.8%) even have more than 100 counts. Thresholds can be applied for minimal expression count per ROI (default ≥ 1). Count data was log-transformed to visualize expression gradients at the single-cell level. A shiny app (SPACE tool) was developed to browse and visualize the expression data: https://www.psb.ugent.be/shiny/space_tool/.

Cell type analysis and clustering were performed using Scanpy (Wolf et al., 2018). For quality control, cells with less than 5 gene counts were filtered out. To better distinguish expression patterns, cells with less than 3 genes expressed were filtered out. Dimension reduction method principal component analysis (PCA) and uniform manifold approximation and projection (UMAP) (McInnes et al., 2018) were performed with different settings (dimensions = 1:5 and 1:9, number of nearest neighbors = 50 or 200 and resolution = 0.1 and 0.5). PCA analysis and projection on the section are available in the SPACE tool.

Pairwise expression analysis was done by either direct comparison (counts of gene 1 ≥ 1 and counts gene 2 ≥ 1) or relative comparison (counts gene 1 / (counts gene 1 + counts gene 2)). Visualization of the expression profiles and pairwise comparisons at the single-cell level were done using the SPACE tool.

## Data availability

SPACE tool: https://www.psb.ugent.be/shiny/space_tool/

**Username:** reviewer

**Password:** 6N89QKTL

R scripts/packages:

Image correction: https://github.com/peng-lab/BaSiCPy

Segmentation model created using https://github.com/BIOP/qupath-extension-cellpose

---

**Supplemental figure S1:**
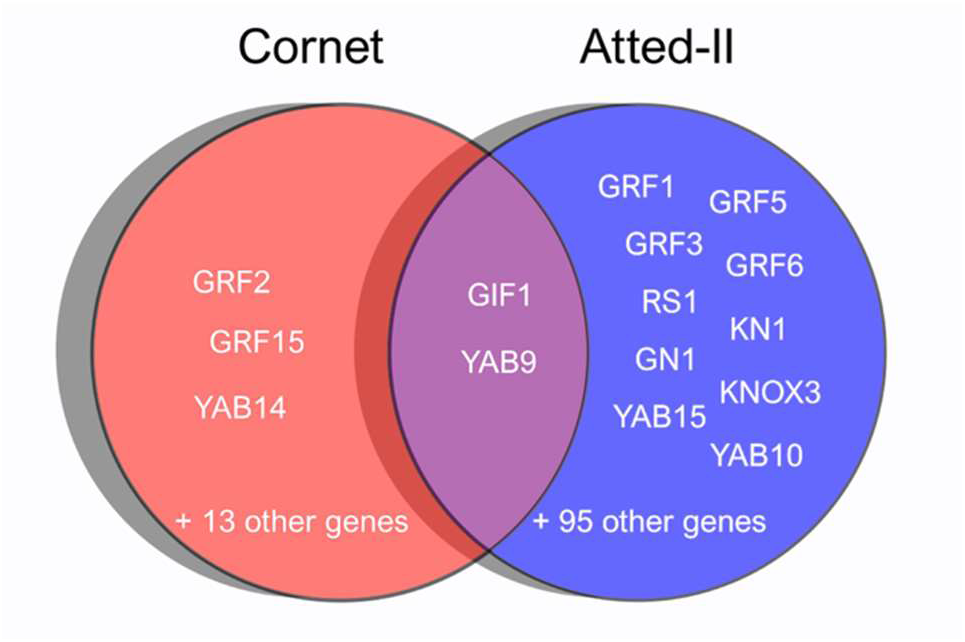
Venn diagram of two co-expression analysis with *PLA1* using CORNET (left) and ATTED-II (right) on bulk RNAseq datasets. Genes identified using both methods are shown in the overlap. Full list of co-expressed genes can be found in **supplemental table 1 and 2**.

**Supplemental figure S2:**
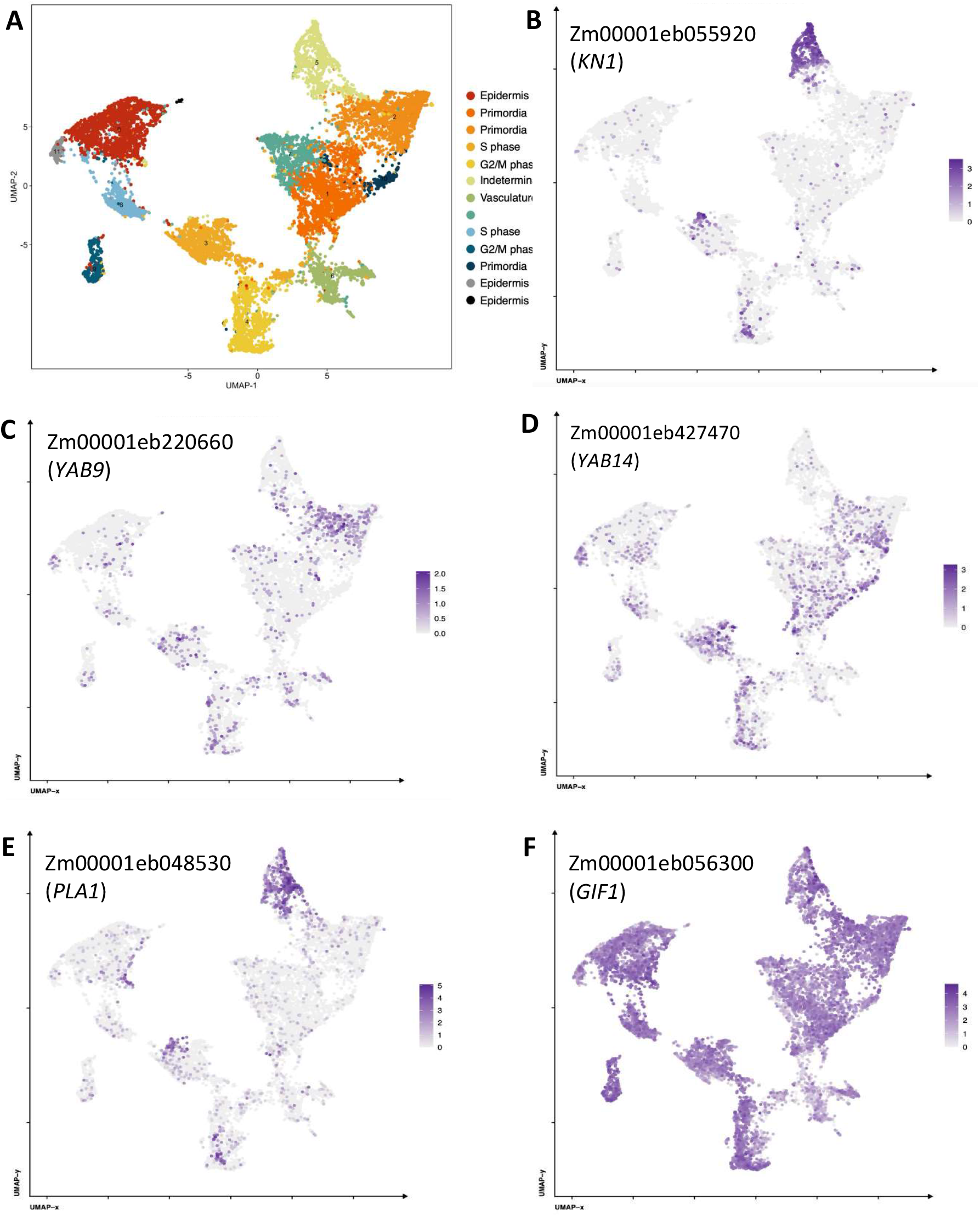
Cell type specific expression of *KN1, YAB9, YAB14, PLA1* and *GIF1* in the maize shoot apex (SAM + P6, Satterlee et. al, 2020). (**A**) UMAP and cell classification of the SAM + P6 dataset. Cell type specific expression of (**B**) *KN1*, (**C**) *YAB9*, (**D**) *YAB14*, (**E**) *PLA1* and (**F**) *GIF1*.

**Supplemental figure S3:**
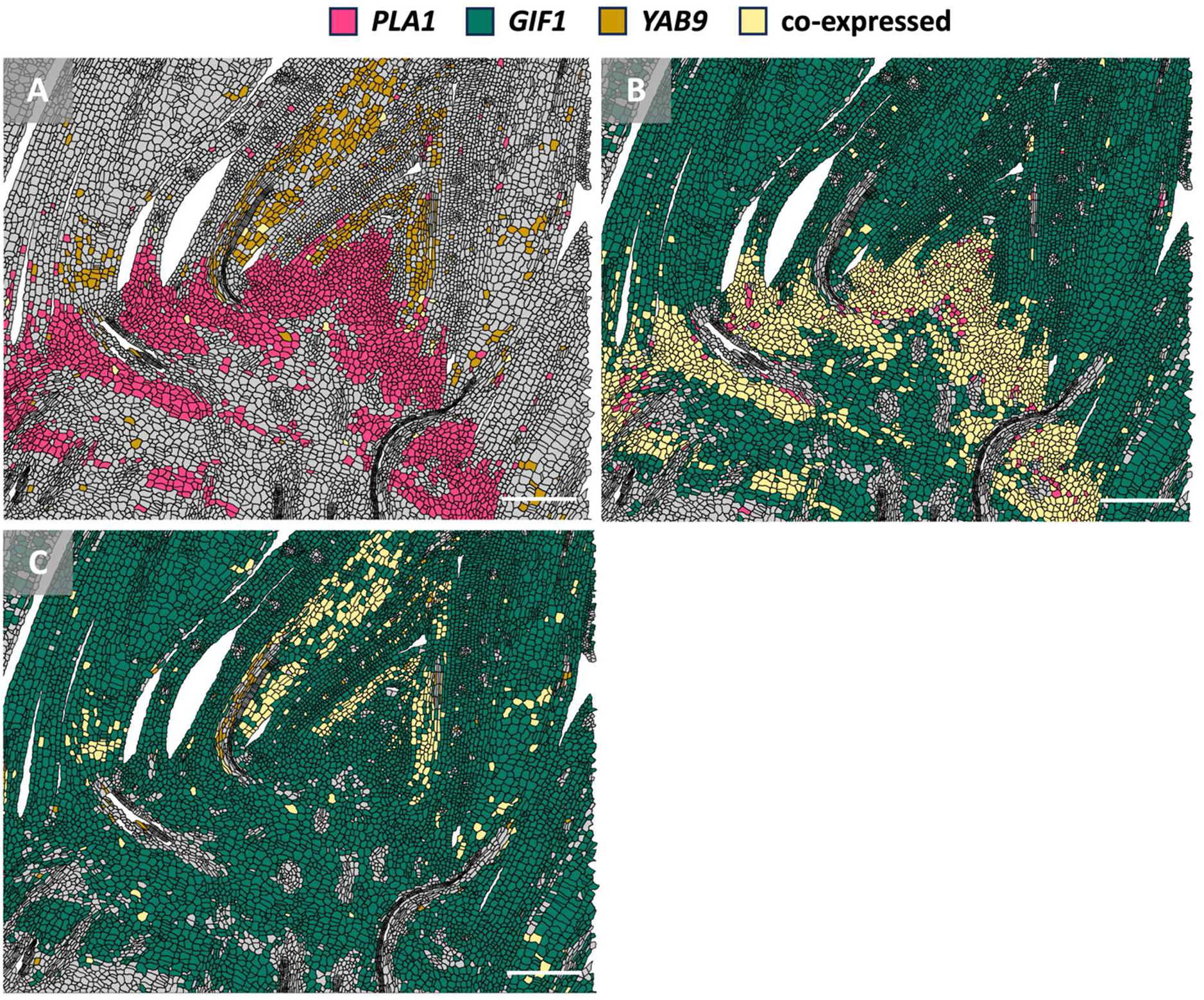
Co-expression of *PLA1, GIF1* and *YAB9* in the maize shoot apex. Pairwise comparison between (**A**) *PLA1* and *YAB9*, (**B**) *PLA1* and *GIF1* and (**C**) *GIF1* and *YAB9*. Scale bars = 200 µm. Molecular cartography was used to generate the spatial maps.

**Supplemental figure S4:**
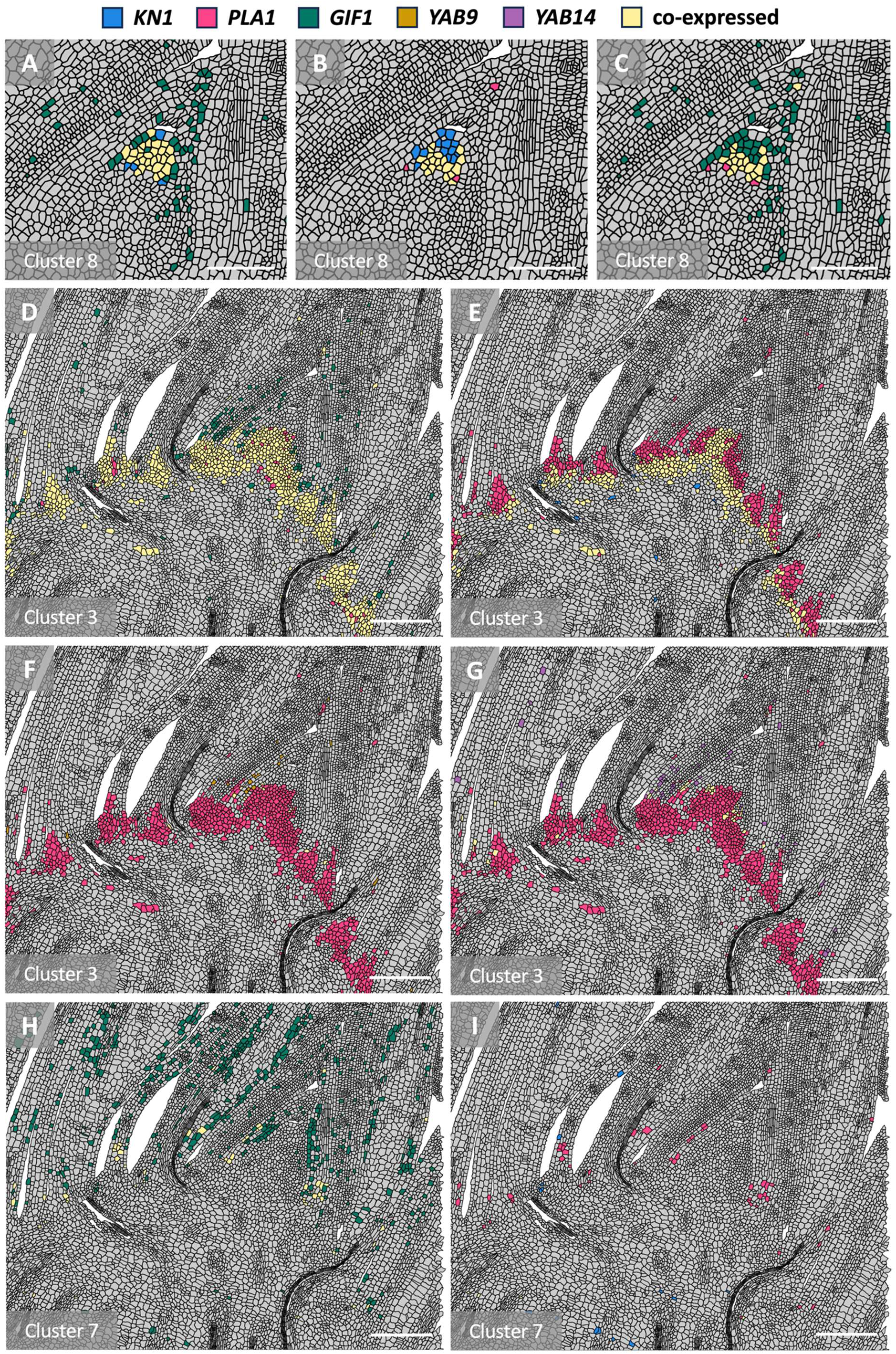
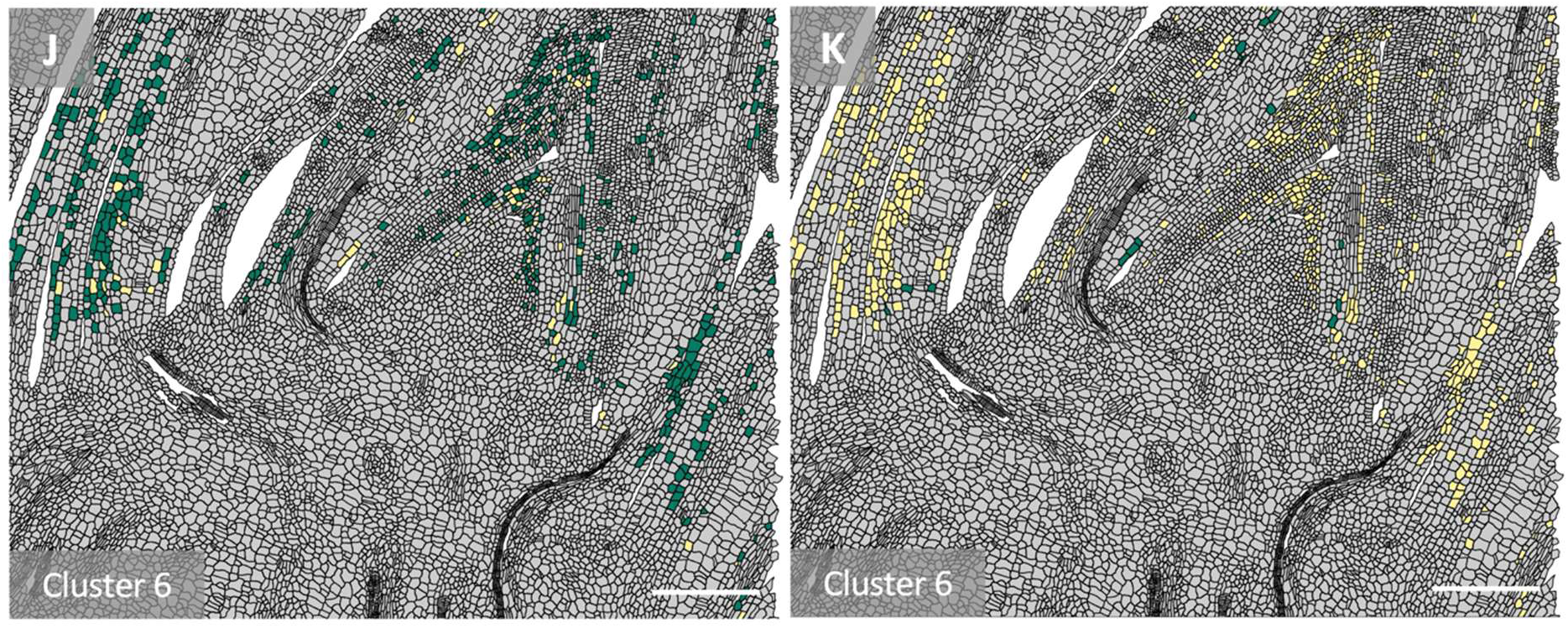
Cell type specific co-expression of *KN1, PLA1, GIF1, YAB9* and *YAB14* in the maize shoot apex. (**A-C**) Cluster 8 (tip of the SAM): pairwise comparison between (**A**) *KN1* and *GIF1,* (**B**) *KN1* and *PLA1,* (**C**) *PLA1* and *GIF1.* Scale bars = 100 µm. (**D-G**) Cluster 3 (boundary): pairwise comparison between (**D**) *PLA1* and *GIF1,* (**E**) *PLA1* and *KN1,* (**F**) *PLA1* and *YAB9* and (**G**) *PLA1* and *YAB14*. Scale bars = 200 µm. (**H-I**) Cluster 7 (proximal part of the base of the leaf): pairwise comparison between (**H**) *PLA1* and *GIF1* and (**I**) *PLA1* and *KN1*. Scale bars = 200 µm. (**J-K**) Cluster 7 (distal part of the base of the leaf): pairwise comparison between (**J**) *GIF1* and *YAB9* and (**K**) *GIF1* and *YAB14*. Scale bars = 200 µm. Molecular cartography was used to generate the spatial maps.

**Supplemental figure S5:**
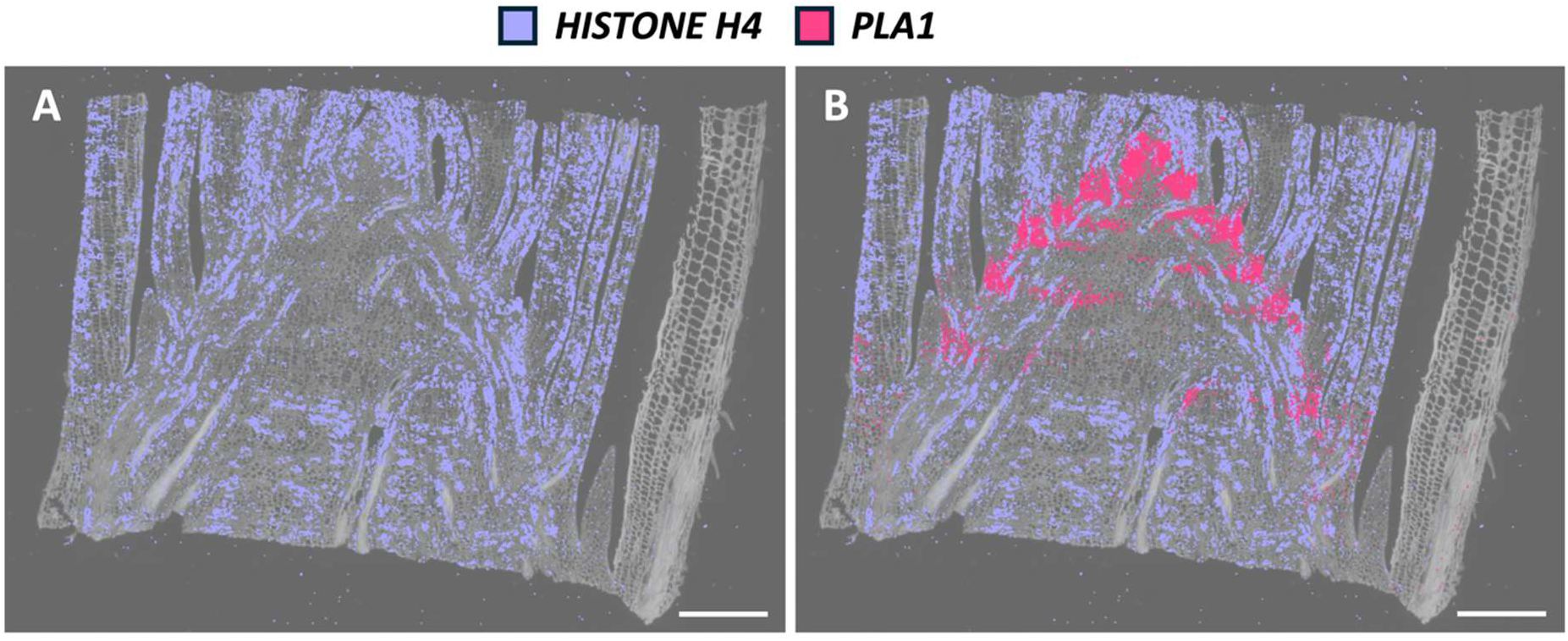
Expression of *HISTONE H4* and *PLA1* in the maize shoot apex (Laureyns et al., 2022). (**A**) Expression of *HISTONE H4* and (**B**) co-expression of *PLA1* with *HISTONE H4*. Scale bars = 100 µm. *In situ* sequencing was used to generate the spatial maps.

**Supplemental figure S6:**
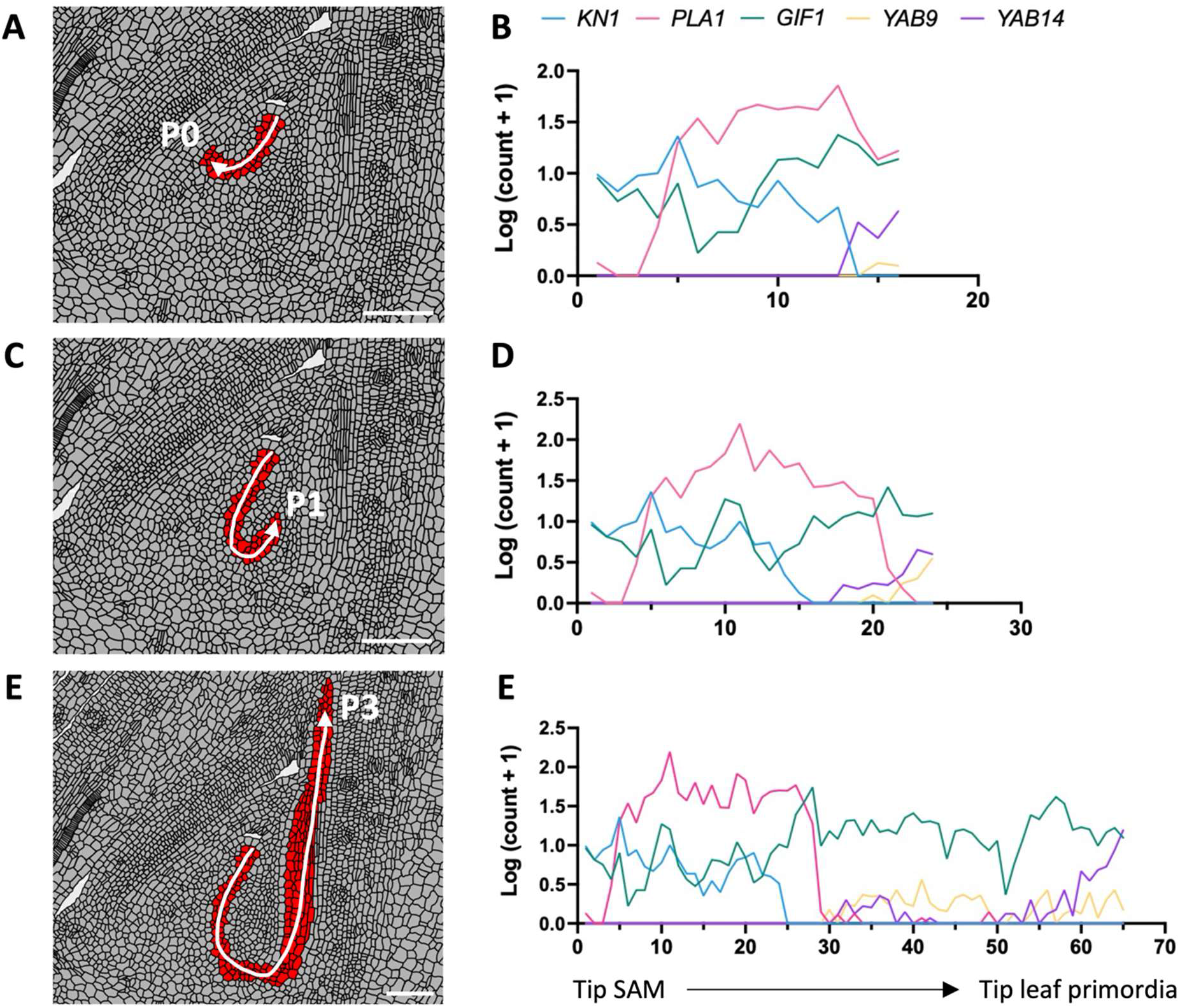
Transcriptional gradients of *KN1, PLA1, GIF1, YAB9* and *YAB14* along a predefined spatial trajectory in the maize shoot apex. Selected cells (red) of the predefined spatial trajectory going from the tip of the SAM to (**A-B**) P0, (**C-D**) P1 and (**D-E**) P3. Transcript counts per cell were log transformed and the average of 3-5 horizontally aligned cells are shown across the spatial trajectory. Molecular cartography was used to generate the spatial maps.

**Supplemental figure S7:**
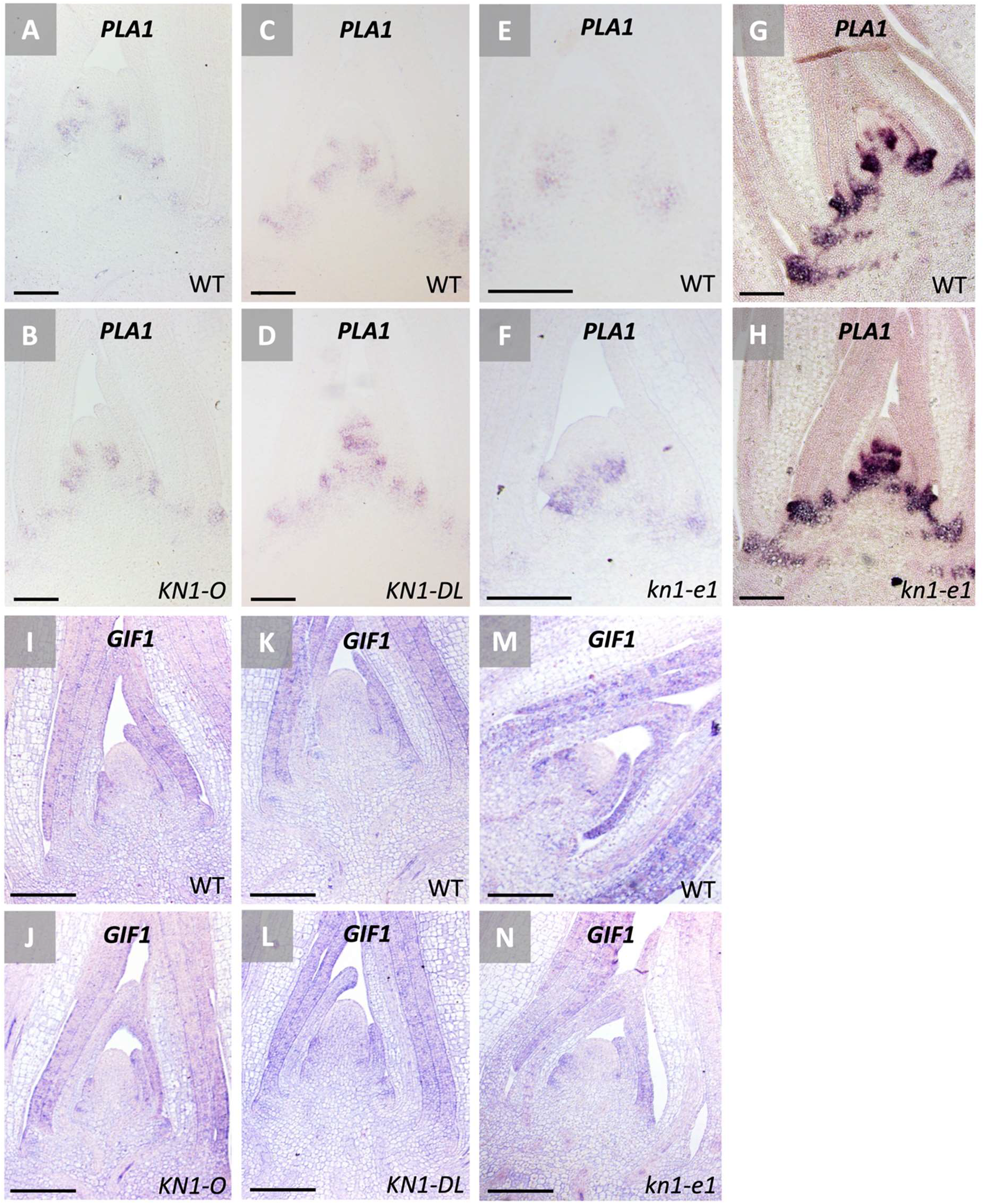
*In situ* hybridization (ISH) of *PLA1* and *GIF1* in the maize shoot apex of *KN1* perturbations. ISH of (**A-H**) *PLA1* and (**I-N**) *GIF1* in the (**A-B-I-J**) *KN1-O* mutant, (**C-D-K-L**) *KN1-DL* mutant, (**E-F-M-N**) *kn1-e1* mutant, (**G-H**) and the permissive *kn1-e1* background and their respective wild type (WT) control. Scale bars = 100 µm.

**Supplemental figure S8:**
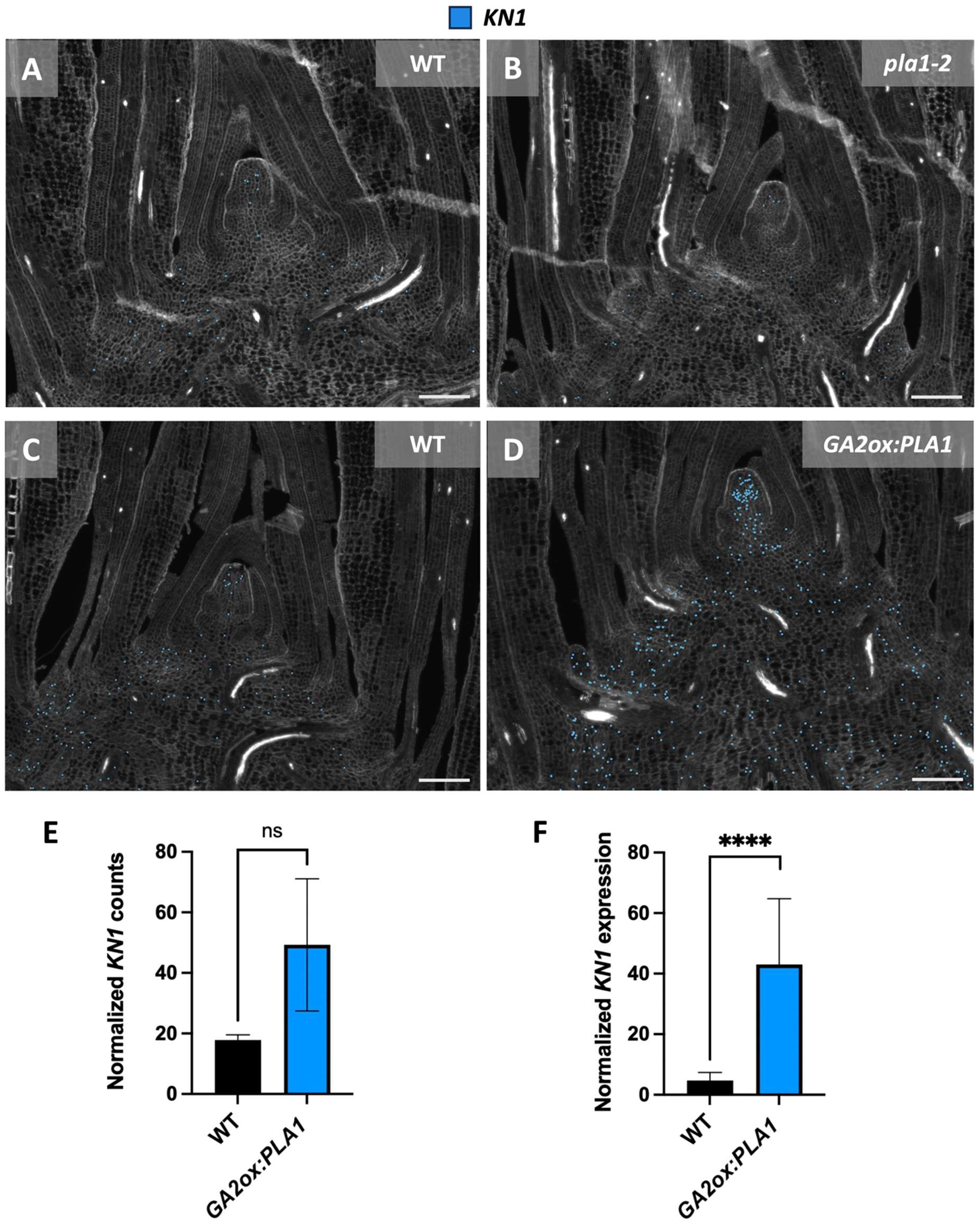
*KN1* expression in the maize shoot apex of the *pla1-2* mutant and the *GA2ox:PLA1* line. *KN1* expression in (**A**) wild type (WT) and (**B**) the *pla1-2* mutant. *KN1* expression in (**C**) wild type and (**D**) the *GA2ox:PLA1* line. (**E**) Normalized counts of *KN1* in *GA2ox:PLA1* compared with wild type using *in situ* sequencing. Significance was calculated by a Welch’s t test (n = 2). ns = non-significant. Scale bars = 100 µm. *In situ* sequencing was used to generate the spatial maps. (**F**) Normalized *KN1* expression in a transcriptome analysis of the basal half cm of leaf 4 from *GA2ox:PLA1* plants compared with wild type (Sun et al., 2017). For statistical analysis see Sun et al., 2017. **** = P ≤ 0.0001.

**Supplemental figure S9:**
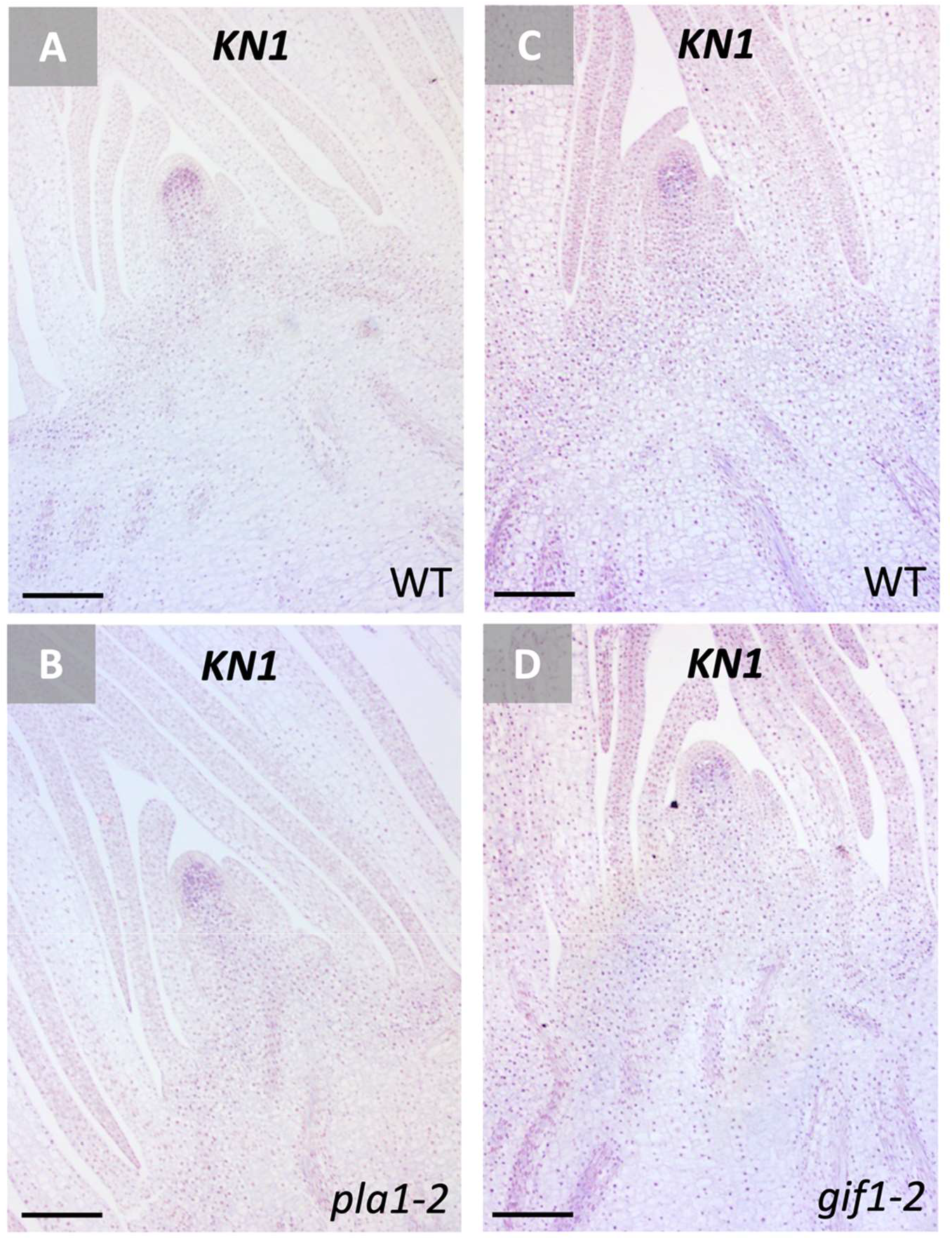
*In situ* hybridization (ISH) of *KN1* in the maize shoot apex of the *pla1-2* mutant and the *gif1-2* mutant. ISH of *KN1* in (**A**) wild type and (**B**) the *pla1-2* mutant, (**C**) wild type and (**D**) the *gif1-2* mutant. Scale bars = 100 µm.

**Supplemental figure S10:**
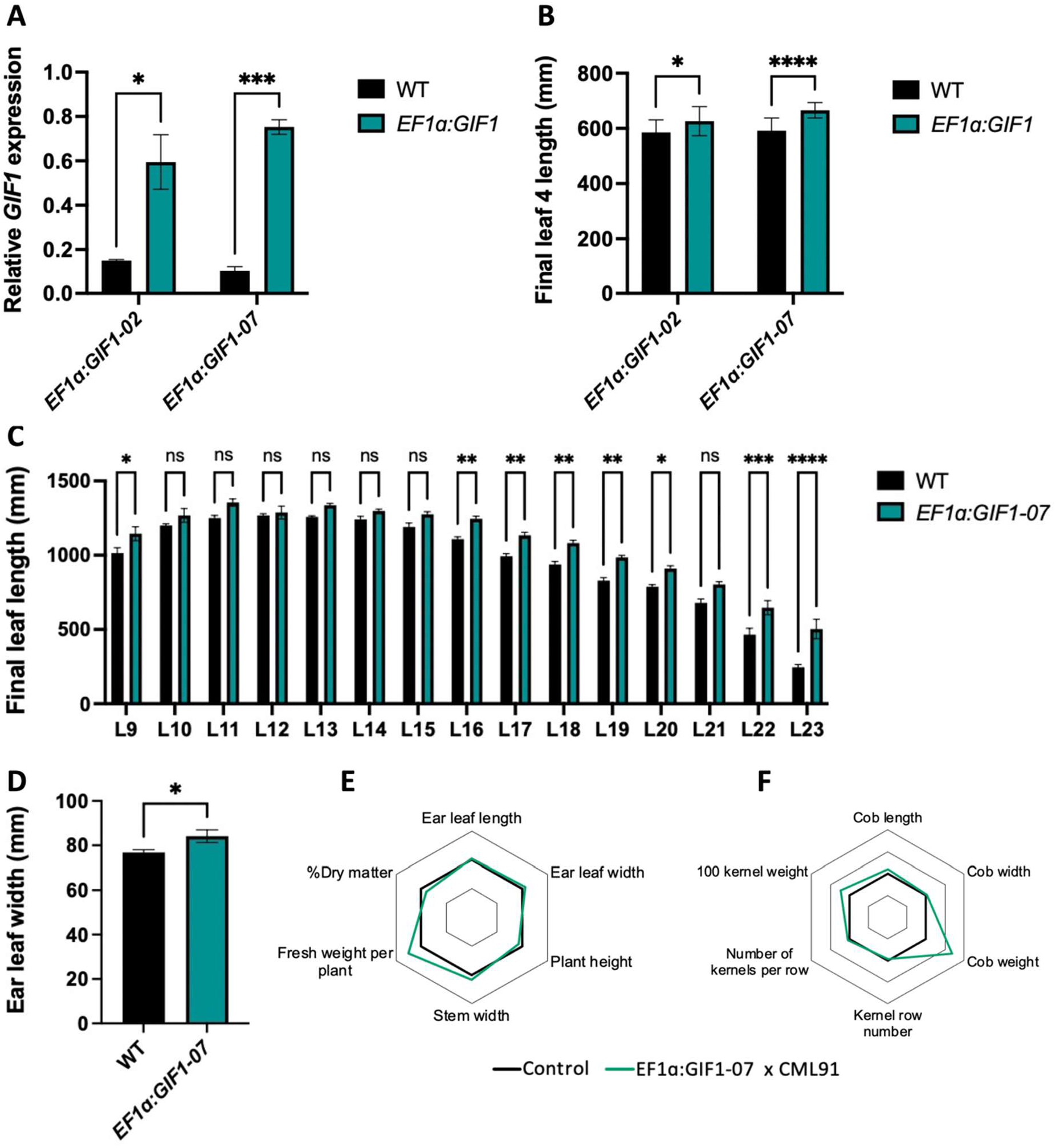
High overexpression of *GIF1* increases growth in growth chamber, greenhouse and field conditions. (**A**) *GIF1* expression and (**B**) final leaf 4 length in the *GIF1* overexpression lines driven by the *EF1α* promotor (*EF1α:GIF1-02* and *EF1α:GIF1-07*). Significance was calculated by a two-way analysis of variance with Bonferroni correction (n ≥ 3). (**C-D**) Phenotypes for (**C**) final leaf length and (**D**) ear leaf width in the greenhouse. Significant differences for final leaf length were calculated by a two-way analysis of variance with Bonferroni correction (n ≥5). Significant differences for ear leaf width were calculated by a Welch’s t test (n ≥ 5). (**E-F**) Growth related phenotypes of *EF1ɑ:GIF1-07* x CML91 hybrids in field conditions. Statistical analysis can be found in **Supplemental table T7**. * = P ≤ 0.05, ** = P ≤ 0.01, *** = P ≤ 0.001 and **** = P ≤ 0.0001. ns = non-significant.

**Supplemental figure S11:**
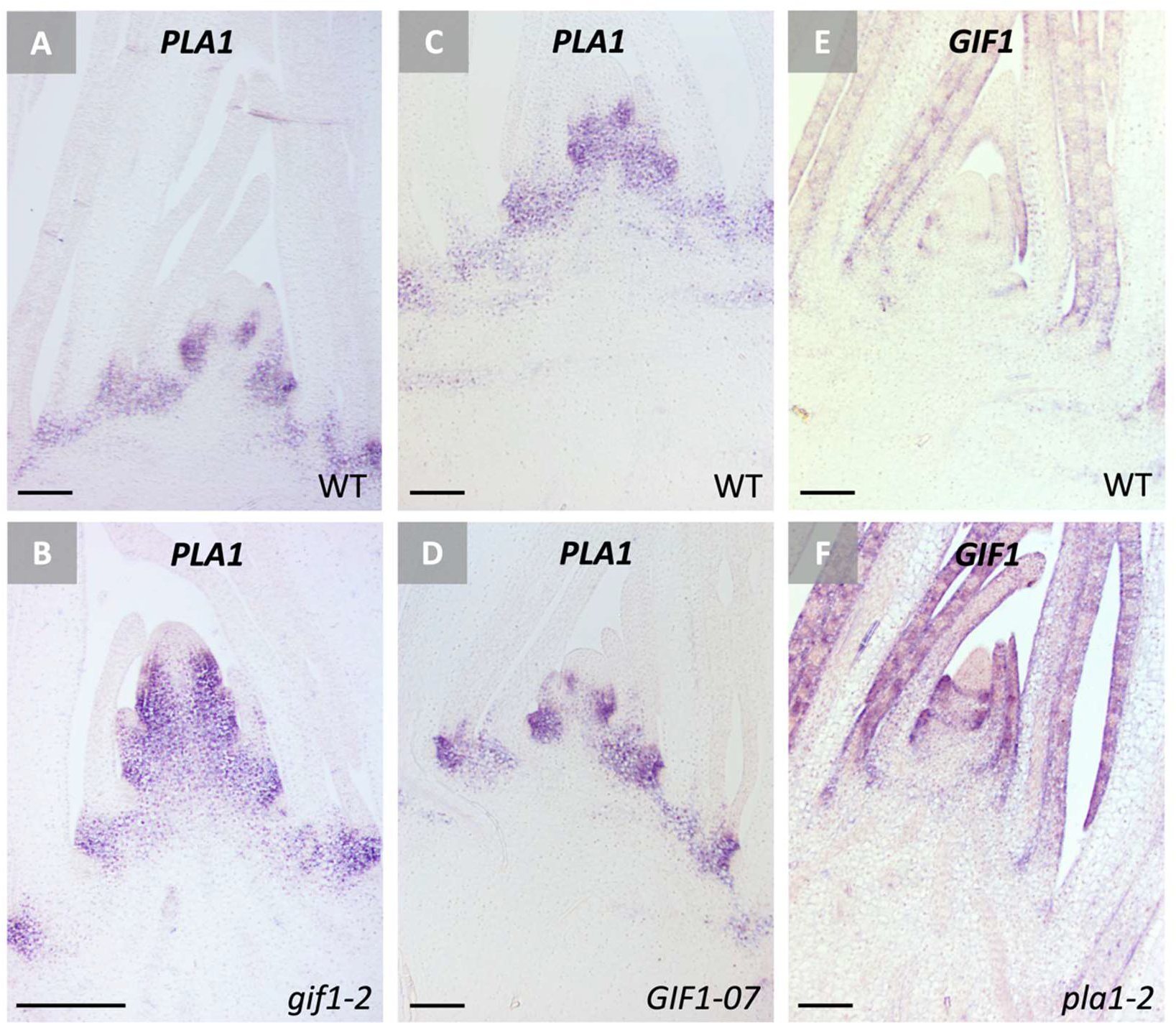
*In situ* hybridization (ISH) validation for figure 4D-I. ISH of *PLA1* in (**A**) wild type and (**B**) the *gif1-2* mutant, (**C**) wild type and (**D**) the *EF1ɑ:GIF-07* overexpression line. ISH for *GIF1* in (**E**) wild type and (**F**) the *pla1-2* mutant. Scale bars = 100 µm.

**Supplemental figure S12:**
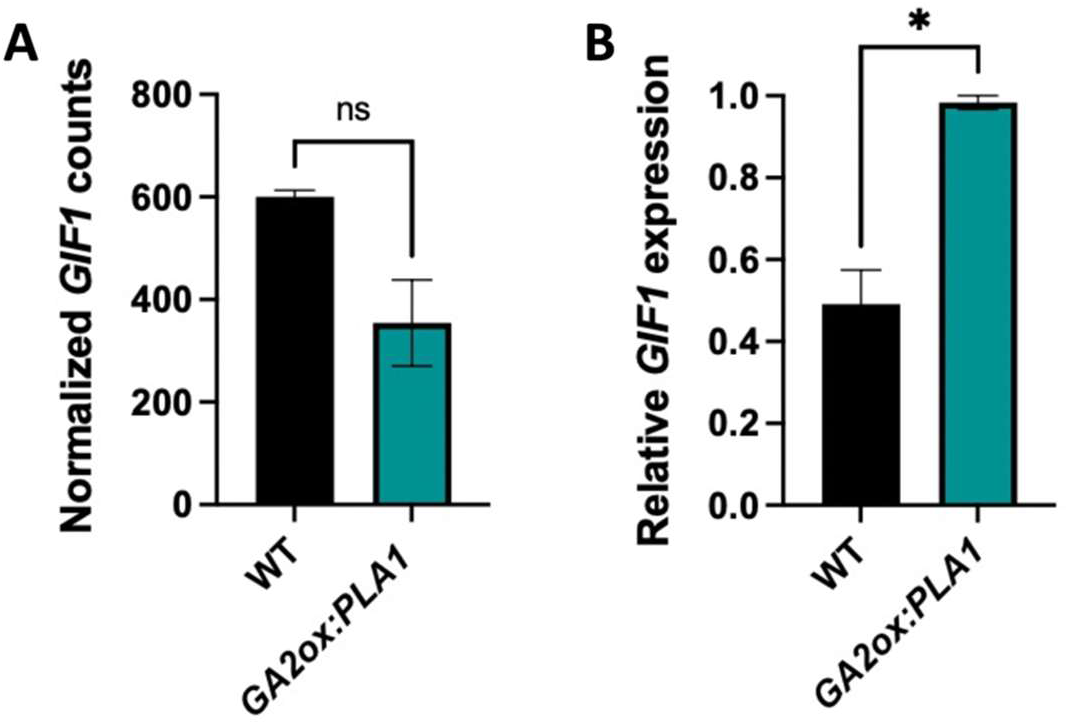
*GIF1* expression in the *GA2ox:PLA1* line. (**A**) Normalized counts of *GIF1* in the shoot apex section of *GA2ox:PLA1* compared with wild type (WT) using *in situ* sequencing. (**B**) Relative expression of *GIF1* in *GA2ox:PLA1* compared with wild type in the leaf 4 division zone, determined by RT-QPCR. Significance was calculated by a Welch’s t test (n ≥ 2). * = P ≤ 0.05. ns = non-significant.

**Supplemental figure S13:**
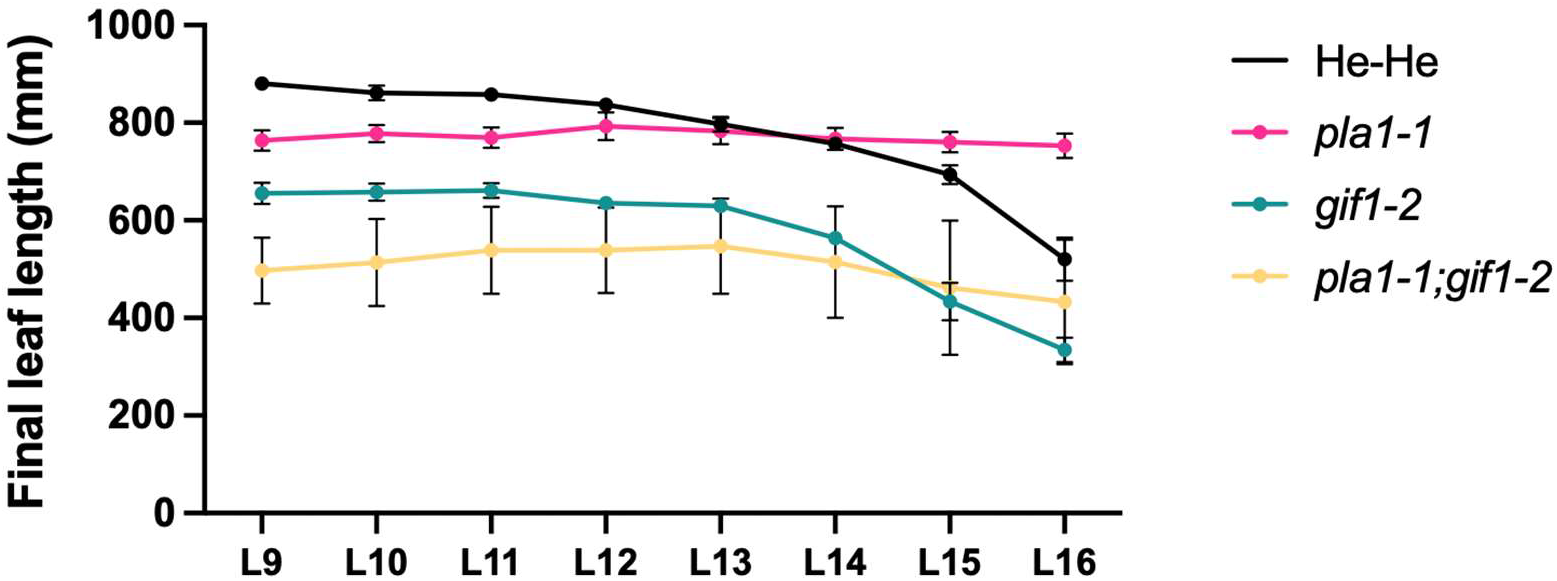
Final leaf lengths of the *pla1-1;gif1-2* double mutants and segregating single *pla1-1* and *gif1-2* mutants in the greenhouse. Statistical analysis can be found in **supplementary table T8**.

**Supplemental figure S14:**
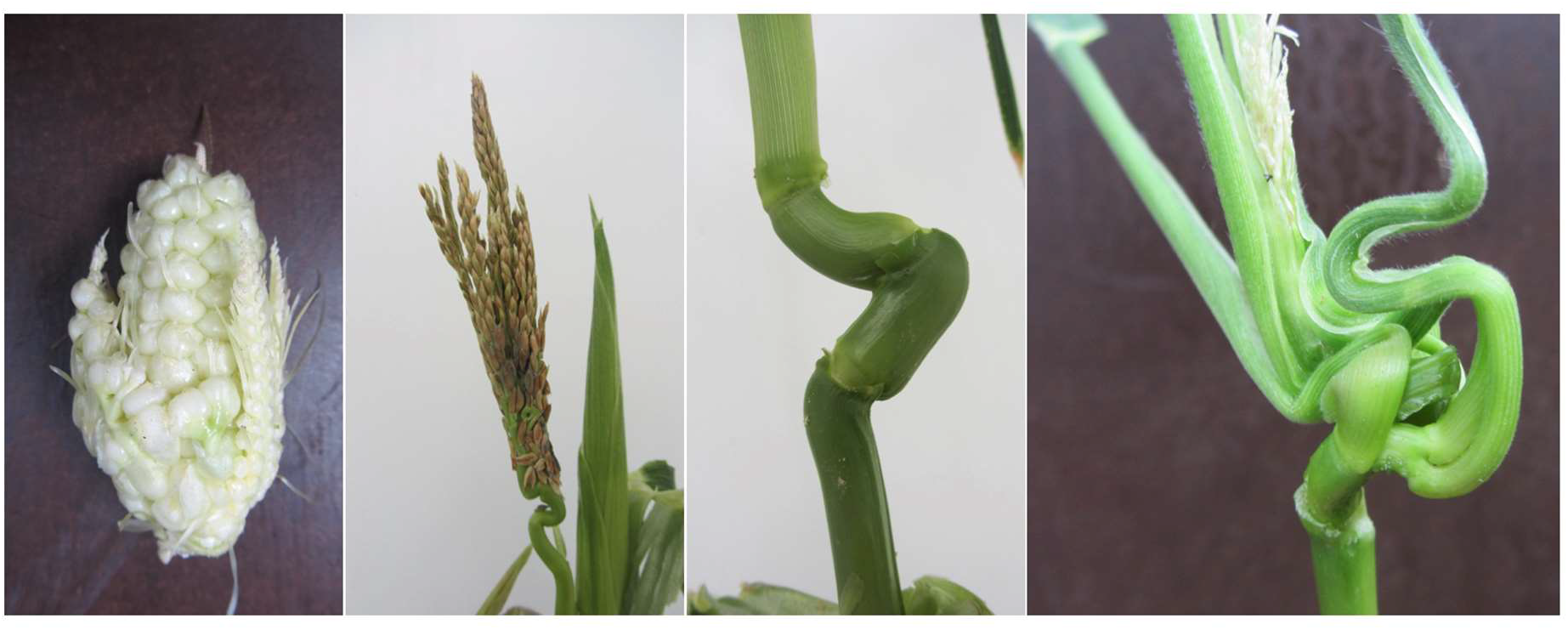
Phenotypes of *pla1-1;gif1-2* double mutants. Double mutants could not be propagated as they mostly did not develop functional reproductive organs. Phenotypes observed included fasciated, branched ears and asymmetric internode.

**Supplemental figure S15:**
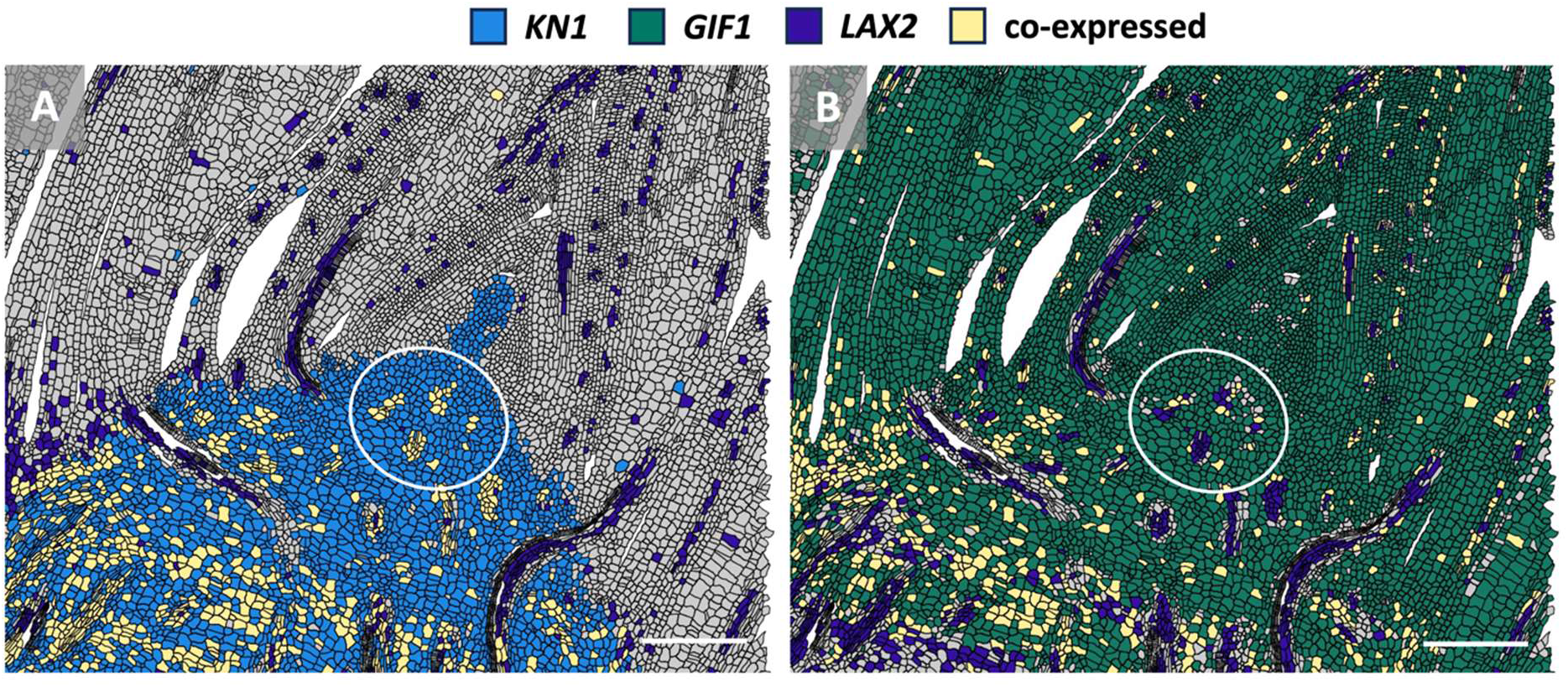
*KN1* and *GIF1* are co-expressed with *LAX2* in developing provascular cells in the ribzone of the shoot apex (white circle). Pairwise comparison of (**A**) *KN1* and *LAX2* and (**B**) *GIF1* and *LAX2*. Scale bars = 200 µm. Molecular cartography was used to generate the spatial maps.

**Supplemental figure S16:**
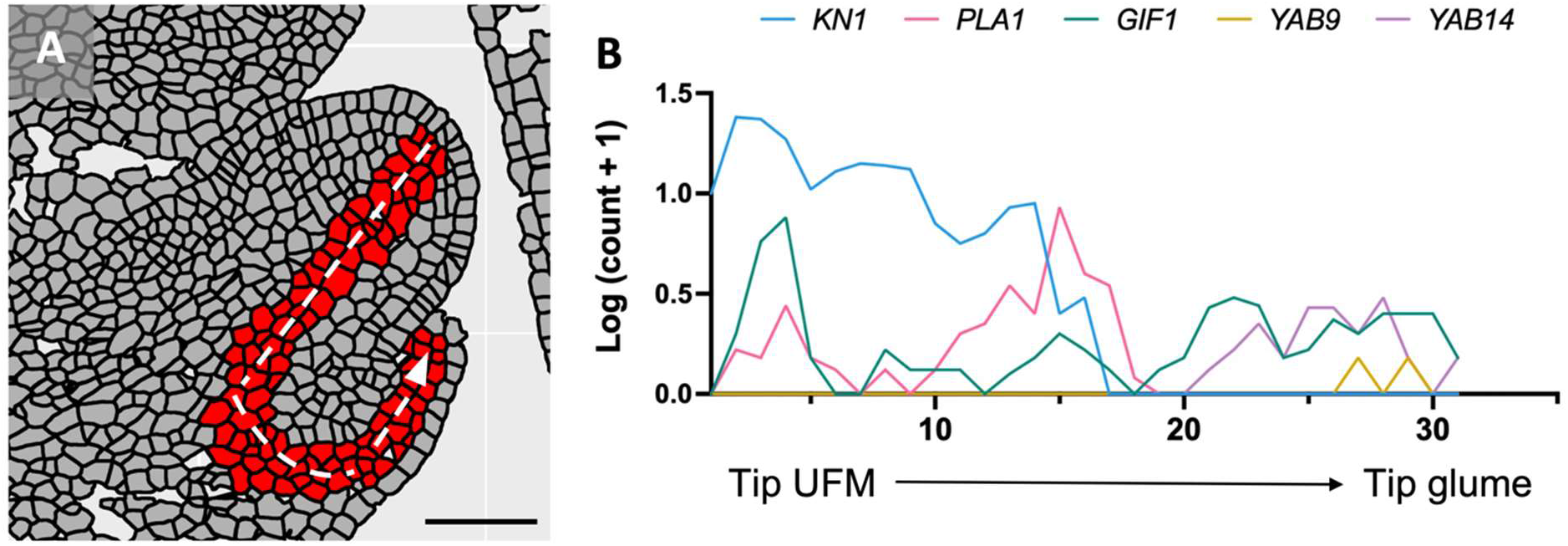
Transcriptional gradients of *KN1, PLA1, GIF1, YAB9* and *YAB14* along a predefined spatial trajectory in a young ear spikelet. (**A**) Selected cells (red) for the predefined spatial trajectory going from the tip of the upper floral meristem (UFM) to the tip of the glume (white dashed arrow). Scale bar = 50 µm. (**B**) Transcript counts per cell were log-transformed and the average of 3-5 horizontally aligned cells are shown across the spatial trajectory.

**Supplemental table T1:**
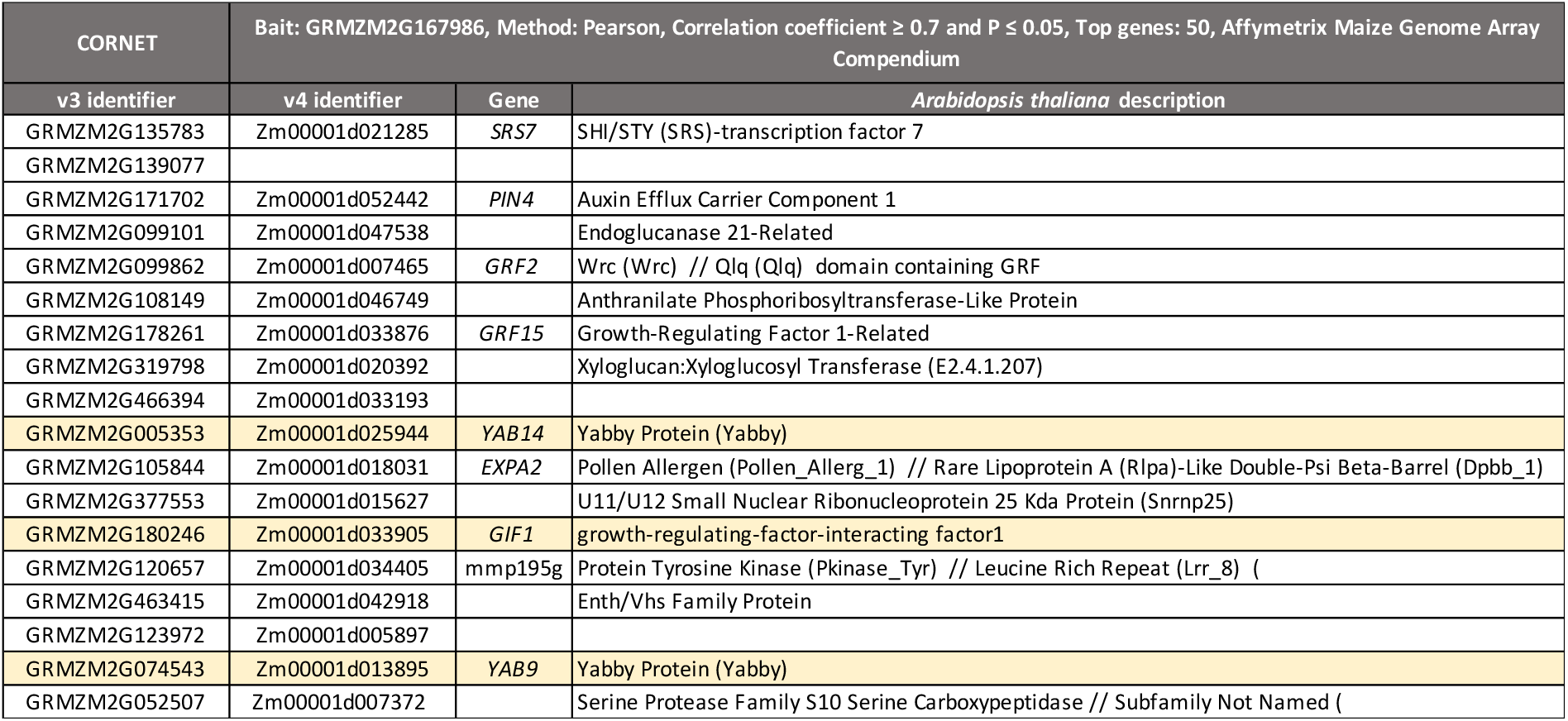
Genes identified to be co-expressed with PLA1 using CORNET (correlation coefficient ≥ 0.7 and P ≤ 0.05). Genes marked in yellow are in the molecular cartography dataset.

**Supplemental table T2:** Genes identified to be co-expressed with PLA1 using Atted-II (P ≤ 0.01). Genes marked in yellow are in the molecular cartography dataset.

| Gene |  | Name/function | P-value |
| --- | --- | --- | --- |
| v3 identifier | v4 identifier |  |  |
| GRMZM2G017087 | Zm00001d033859 | knotted 1 | 1.00E-03 |
| GRMZM2G168371 | Zm00001d018433 | putative protein of unknown function (DUF640) domain family | 1.00E-03 |
| GRMZM2G163724 | Zm00001d005298 | uncharacterized LOC100383466 | 1.00E-03 |
| GRMZM2G346920 | Zm00001d031840 | uncharacterized LOC100283050 | 1.00E-03 |
| GRMZM2G359779 | Zm00001d004094 | protein SMAX1-LIKE 4 | 1.00E-03 |
| GRMZM2G129413 | Zm00001d030310 | auxin import carrier 1 | 1.00E-03 |
| GRMZM2G032547 | Zm00001d025136 | Protein SMAX1-LIKE 4 | 1.00E-03 |
| GRMZM2G123972 | Zm00001d005897 | uncharacterized LOC100276018 | 1.00E-03 |
| GRMZM2G028151 | Zm00001d034204 | uncharacterized LOC100217129 | 1.00E-03 |
| GRMZM2G146688 | Zm00001d007840 | uncharacterized LOC100502174 | 1.00E-03 |
| GRMZM2G124566 | Zm00001d051456 | uncharacterized LOC100286247 | 1.00E-03 |
| GRMZM2G455642 | Zm00001d037338 | uncharacterized LOC100216867 | 1.00E-03 |
| GRMZM2G034876 | Zm00001d017742 | growth-regulating factor 1-like (GRF1) | 1.00E-03 |
| GRMZM2G108865 | Zm00001d034163 | uncharacterized LOC100277644 | 1.00E-03 |
| GRMZM2G103983 | Zm00001d032610 | tonoplast intrinsic protein 3 | 1.00E-03 |
| GRMZM2G025685 | Zm00001d022295 | uncharacterized LOC100277279 | 1.00E-03 |
| GRMZM2G305446 | Zm00001d013956 | tonoplast intrinsic protein 3 | 1.00E-03 |
| GRMZM2G040517 | Zm00001d038741 | uncharacterized LOC100193845 | 1.00E-03 |
| AC210003.2_FG004 | Zm00001d002004 | uncharacterized LOC100274427 | 1.00E-03 |
| GRMZM2G105335 | Zm00001d052112 | growth-regulating factor 3-like (GRF3) | 1.00E-03 |
| GRMZM2G529859 | Zm00001d017391 | protein YABBY 4 (YAB15) | 1.00E-03 |
| GRMZM2G337229 | Zm00001d002768 | oleosin 1 | 1.00E-03 |
| GRMZM2G032865 | Zm00001d038239 | uncharacterized LOC100278467 | 1.00E-03 |
| GRMZM2G052507 | Zm00001d007372 | serine carboxypeptidase-like 45 | 1.00E-03 |
| GRMZM2G025793 | Zm00001d034003 | late embryogenesis abundant protein 47 | 1.00E-03 |
| GRMZM2G180246 | Zm00001d033905 | growth-regulating-factor-interacting factor1 (*GIF1*) | 1.00E-03 |
| GRMZM2G008234 | Zm00001d018731 | uncharacterized LOC100502464 | 1.00E-03 |
| GRMZM2G024898 | Zm00001d002452 | probable WRKY transcription factor 14 | 1.00E-03 |
| GRMZM2G168588 | Zm00001d021447 | LysM domain-containing GPI-anchored protein 2 | 1.00E-03 |
| GRMZM2G022606 | Zm00001d023246 | BTB/POZ domain and ankyrin repeat-containing protein NH5.2 | 1.00E-03 |
| GRMZM2G480903 | Zm00001d003927 | Grx_C8 - glutaredoxin subgroup III | 1.00E-03 |
| GRMZM2G133398 | Zm00001d042396 | viviparous 1 | 1.00E-03 |
| GRMZM2G033566 | Zm00001d046254 | anther-specific proline-rich protein APG | 1.00E-03 |
| GRMZM2G052111 | Zm00001d039218 | putative leucine-rich repeat receptor-like protein kinase family protein | 1.00E-03 |
| GRMZM5G819222 | Zm00001d021063 | uncharacterized LOC100273006 | 1.00E-03 |
| GRMZM2G114276 | Zm00001d002599 | putative leucine-rich repeat receptor-like protein kinase family protein | 1.00E-03 |
| GRMZM2G074543 | Zm00001d013895 | yabby 9 | 1.00E-02 |
| GRMZM2G171702 | Zm00001d052442 | uncharacterized LOC100285745 | 1.00E-02 |
| GRMZM2G028041 | Zm00001d018742 | rough sheath 1 | 1.00E-02 |
| GRMZM2G452178 | Zm00001d007842 | homeobox protein *gnarley 1* | 1.00E-02 |
| GRMZM2G082586 | Zm00001d020790 | uncharacterized LOC100280340 | 1.00E-02 |
| GRMZM2G108798 | Zm00001d036426 | protein SHI RELATED SEQUENCE 1 | 1.00E-02 |
| GRMZM2G017882 | Zm00001d015394 | uncharacterized LOC103626673 | 1.00E-02 |
| GRMZM2G393433 | Zm00001d035266 | uncharacterized LOC103629107 (*cuc2*) | 1.00E-02 |
| GRMZM2G025685 | Zm00001d022524 | WUSCHEL-related homeobox 11 | 1.00E-02 |
| GRMZM2G094241 | Zm00001d010948 | liguleless 4 | 1.00E-02 |
| AC205471.4_FG008 | Zm00001d010813 | leucine-rich repeat receptor-like kinase protein FLORAL ORGAN | 1.00E-02 |
| GRMZM2G087741 | Zm00001d040611 | liguleless 3 | 1.00E-02 |
| GRMZM2G015666 | Zm00001d005939 | uncharacterized LOC100285428 | 1.00E-02 |
| GRMZM2G140739 | Zm00001d017789 | uncharacterized LOC100277409 | 1.00E-02 |
| GRMZM2G166413 | Zm00001d011757 | uncharacterized LOC100192730 | 1.00E-02 |
| GRMZM2G061515 | Zm00001d043244 | probable indole-3-acetic acid-amido synthetase GH3.1 | 1.00E-02 |
| GRMZM2G000743 | Zm00001d033861 | KNOX1 domain containing protein (KNOX3) | 1.00E-02 |
| GRMZM2G358386 | Zm00001d021705 | uncharacterized LOC100275254 | 1.00E-02 |
| GRMZM5G832409 | Zm00001d035439 | knotted related homeobox 5 | 1.00E-02 |
| AC233943.1_FG002 | Zm00001d039694 | ramosa 2 | 1.00E-02 |
| GRMZM2G132077 | Zm00001d030698 | uncharacterized LOC100276911 | 1.00E-02 |
| GRMZM2G097683 | Zm00001d014762 | protein SHI RELATED SEQUENCE 1 | 1.00E-02 |
| GRMZM2G089619 | Zm00001d003645 | ZF-HD homeobox protein | 1.00E-02 |
| GRMZM2G138750 | Zm00001d039650 | putative cytochrome P450 superfamily protein | 1.00E-02 |
| GRMZM2G128807 | Zm00001d014814 | uncharacterized LOC100274388 | 1.00E-02 |
| GRMZM2G003424 | Zm00001d029010 | uncharacterized LOC100384253 | 1.00E-02 |
| GRMZM2G151955 | Zm00001d032142 | probable LRR receptor-like serine/threonine-protein kinase At2g23950 | 1.00E-02 |
| GRMZM2G069365 | Zm00001d051573 | zinc finger homeodomain protein 1 | 1.00E-02 |
| GRMZM2G445684 | Zm00001d031416 | Zinc finger protein WIP2 | 1.00E-02 |
| GRMZM2G463493 | Zm00001d008715 | LRR receptor-like serine/threonine-protein kinase | 1.00E-02 |
| GRMZM2G165428 | Zm00001d011211 | probable inactive leucine-rich repeat receptor kinase XIAO | 1.00E-02 |
| GRMZM2G440785 | Zm00001d012350 | uncharacterized LOC100217015 | 1.00E-02 |
| GRMZM2G113840 | Zm00001d031316 | Phosphatidylinositol/phosphatidylcholine transfer protein SFH13 | 1.00E-02 |
| GRMZM2G098813 | Zm00001d026231 | zea floricaula/leafy 1 | 1.00E-02 |
| GRMZM2G378580 | Zm00001d049295 | auxin response factor | 1.00E-02 |
| GRMZM2G068330 | Zm00001d049000 | zinc-finger homeodomain protein 4 | 1.00E-02 |
| GRMZM2G167824 | Zm00001d032502 | yabby homolog10 | 1.00E-02 |
| GRMZM2G060723 | Zm00001d011878 | uncharacterized LOC100272726 | 1.00E-02 |
| GRMZM2G314064 | Zm00001d028986 | WUSCHEL-related homeobox 6 | 1.00E-02 |
| GRMZM2G093595 | Zm00001d038001 | ethylene-responsive transcription factor ABI4 | 1.00E-02 |
| GRMZM2G099097 | Zm00001d051932 | uncharacterized LOC100273712 | 1.00E-02 |
| GRMZM2G125976 | Zm00001d038355 | BEL1-like homeodomain protein 9 | 1.00E-02 |
| GRMZM2G124567 | Zm00001d047512 | uncharacterized LOC100194178 | 1.00E-02 |
| GRMZM2G166524 | Zm00001d040130 | uncharacterized LOC100216564 | 1.00E-02 |
| GRMZM2G039867 | Zm00001d042111 | BTB/POZ domain and ankyrin repeat-containing protein NPR5 | 1.00E-02 |
| GRMZM2G069389 | Zm00001d026591 | uncharacterized LOC100276102 | 1.00E-02 |
| GRMZM2G456512 | Zm00001d038223 | Bzip transcription factor-like | 1.00E-02 |
| GRMZM5G804575 | Zm00001d031913 | uncharacterized LOC100283745 | 1.00E-02 |
| GRMZM2G129147 | Zm00001d026240 | growth-regulating factor 5-like (GRF5) | 1.00E-02 |
| GRMZM2G003784 | Zm00001d002833 | pepsin A | 1.00E-02 |
| GRMZM2G139324 | Zm00001d030834 | uncharacterized LOC100279086 | 1.00E-02 |
| GRMZM2G125832 | Zm00001d041004 | uncharacterized LOC100381796 | 1.00E-02 |
| AC233850.1_FG009 | Zm00001d016735 | Protein SMAX1-LIKE 3 | 1.00E-02 |
|  |  | uncharacterized LOC100382909 | 1.00E-02 |
| GRMZM2G115329 | Zm00001d045940 | uncharacterized LOC100275144 | 1.00E-02 |
| GRMZM2G067908 | Zm00001d019976 | uncharacterized LOC100383547 | 1.00E-02 |
| GRMZM2G178014 | Zm00001d024530 | uncharacterized LOC100191431 | 1.00E-02 |
| GRMZM2G041223 | Zm00001d002429 | growth-regulating factor 6-like | 1.00E-02 |
| GRMZM5G874163 | Zm00001d012731 | Auxin response factor 3 | 1.00E-02 |
| GRMZM2G020940 | Zm00001d028998 | uncharacterized LOC100274379 | 1.00E-02 |
| GRMZM2G027302 | Zm00001d051966 | putative protein of unknown function (DUF640) domain family protein | 1.00E-02 |
| GRMZM5G813804 | Zm00001d024537 | AT-hook motif nuclear-localized protein 23 | 1.00E-02 |
| GRMZM2G038152 | Zm00001d009123 | Galactose oxidase/kelch repeat superfamily protein | 1.00E-02 |
| GRMZM2G123202 | Zm00001d002843 | transcription factor MYB41 | 1.00E-02 |
| GRMZM2G168807 | Zm00001d016599 | major ampullate spidroin 2-2 | 1.00E-02 |
| GRMZM2G123094 | Zm00001d031759 | protein indeterminate-domain 16 | 1.00E-02 |
| GRMZM2G034155 | Zm00001d025002 | LRR receptor-like serine/threonine-protein kinase RCH1 | 1.00E-02 |
| GRMZM2G098643 | Zm00001d044812 | probable auxin efflux carrier component 1c | 1.00E-02 |
| GRMZM2G071101 | Zm00001d052883 | Zinc finger protein WIP2 | 1.00E-02 |
| GRMZM2G072868 | Zm00001d019701 | probable LRR receptor-like serine/threonine-protein kinase At4g37250 | 1.00E-02 |
| GRMZM2G313448 | Zm00001d035506 | interactor of constitutive active ROPs 2: chloroplastic | 1.00E-02 |
| GRMZM2G417229 | Zm00001d017784 | uncharacterized LOC100279191 | 1.00E-02 |

**Supplemental table T3:** Genes selected for molecular cartography.

| Gene | Abbreviation | V5 identifier | V4 identifier | V3 identifier | Expected expression | Expression pattern correct? | Publication expression pattern |
| --- | --- | --- | --- | --- | --- | --- | --- |
| *ARGONAUTE18a* | *AGO18a* | Zm00001eb104820 | Zm00001d006351 | GRMZM2G105250 | Tip of the SAM | Yes | Knauer et al., 2019, |
| *GRF-INTERACTING FACTOR1* | *GIF1* | Zm00001eb056300 | Zm00001d033905 | GRMZM2G180246 | Leaf and meristematic cells | Yes | Laureyns et al., 2022 |
| *DWARF11-LIKE* | *D11-LIKE* | Zm00001eb080110 | Zm00001d003349 | GRMZM2G107199 | Boundary | Yes | Johnston et al., 2014 |
| *JACALIN RELATED LECTIN* | *JRL* | Zm00001eb265890 | Zm00001d035562 | GRMZM2G163406 | Mesophyll cells | Yes | Zhang et al., 2007 |
| *ROUGH SHEATH1* | *RS1* | Zm00001eb299420 | Zm00001d018742 | GRMZM2G028041 | Meristematic cells | Yes | Schneeberger et al., 1995 |
| *TREHALOSE-6-PHOSPHATE PHOSPHATASE4* | *TPP4* | Zm00001eb194210 | Zm00001d052227 | GRMZM2G151044 | Boundary | Yes | Laureyns et al., 2022 |
| *ALTERED PHLOEM DEVELOPMENT* | *APL* | Zm00001eb233310 | Zm00001d015407 | GRMZM2G052544 | Phloem | Yes | De Rybel et al., 2016 |
| *BARELY ANY MERISTEM1d* | *BAM1d* | Zm00001eb009640 | Zm00001d028317 | GRMZM2G043584 | Meristematic cells | Yes | Laureyns et al., 2022 |
| *CUP-SHAPED COTYLEDON2* | *CUC2* | Zm00001eb264380 | Zm00001d035266 | GRMZM2G393433 | Boundary marker | Yes | Johnston et al., 2014 |
| *DYNAMIN RELATED PROTEIN4a* | *DRP4a* | Zm00001eb011320 | Zm00001d028495 | GRMZM2G150827 | Tip of the SAM | Yes | Knauer et al., 2019, |
| *KNOTTED1* | *KN1* | Zm00001eb055920 | Zm00001d033859 | GRMZM2G017087 | meristimatic cells | Yes | Jackson et al.,1994 |
| *LIKE AUXIN RESISTANT 2* | *LAX2* | Zm00001eb026490 | Zm00001d030310 | GRMZM2G129413 | Vasculature | Yes | Leiboff et al.,2015 |
| *LONESOME HIGHWAY1* | *LHW1* | Zm00001eb407240 | Zm00001d023479 | GRMZM2G031656 | Vasculature | Yes | De Rybel et al., 2016 |
| *LONESOME HIGHWAY1b* | *LHW1b* | Zm00001eb139520 | Zm00001d041926 | GRMZM2G069408 | Vasculature | Yes | De Rybel et al., 2016 |
| *PIN-FORMED1d* | *PIN1d* | Zm00001eb372180 | Zm00001d044812 | GRMZM2G098643 | Vasculature | Yes | Laureyns et al., 2022 |
| *PLASTOCHRON1* | *PLA1* | Zm00001eb048540 | Zm00001d033020 | GRMZM2G167986 | Boundary | Yes | Laureyns et al., 2022 |
| *TARGET OF MONOPTEROS5* | *TMO5* | Zm00001eb398190 | Zm00001d047878 | GRMZM2G043854 | Xylem | Yes | De Rybel et al., 2016 |
| *TARGET OF MONOPTEROS5-LIKE1* | *TMO5L1* | Zm00001eb041610 | Zm00001d032213 | GRMZM2G107276 | Xylem | Yes | De Rybel et al., 2016 |
| *ZEA ROOT PROTEIN2* | *ZRP2* | Zm00001eb039010 | Zm00001d031909 | GRMZM2G106980 | Primary root and stem | Yes | Held et al., 1997 |
| *YABBY14* | *YAB14* | Zm00001eb427470 | Zm00001d025944 | GRMZM2G005353 | Adaxial leaf | Yes | Juarez et al., 2004 |
| *YABBY9* | *YAB9* | Zm00001eb220660 | Zm00001d013895 | GRMZM2G074543 | Adaxial leaf | Yes | Juarez et al., 2004 |

**Supplemental table T4:**
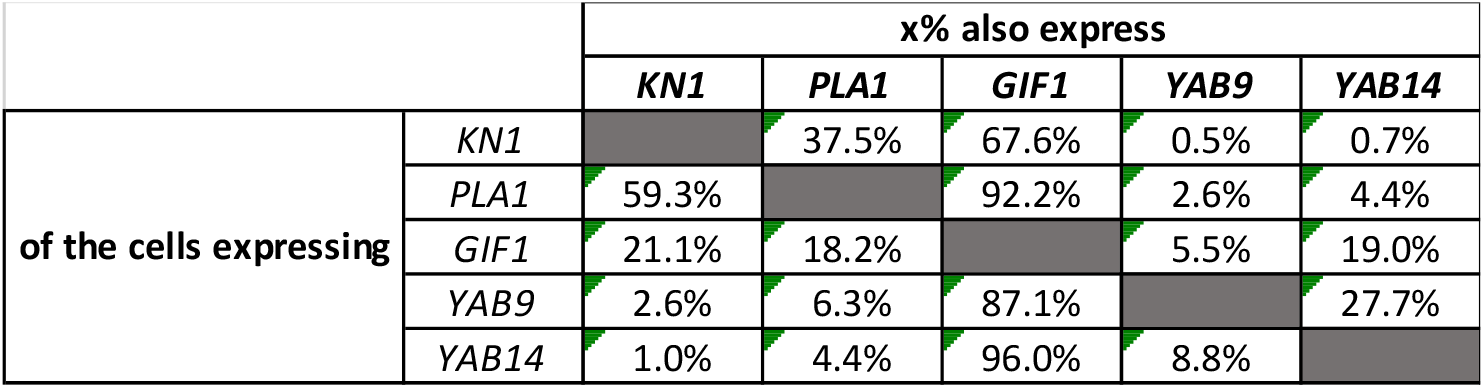
Co-expression in KN1, PLA1, GIF1, YAB9 and YAB14 expressing cells in the maize shoot apex.

**Supplemental table T5:** Growing leaf 4 length over time for a segregating *pla1-2;GA2ox:PLA1* population. Significance was calculated by a mixed model with Bonferroni correction (n ≥ 9). DAL4A = days after leaf 4 appearance. SEM = Standard error of the mean.

| | Wild type<br>Mean ± SEM | *pla1-2*<br>Mean ± SEM | *GA2ox:PLA1*<br>Mean ± SEM | *pla1-2;GA2ox:PLA1*<br>Mean ± SEM |
| --- | --- | --- | --- | --- |
| 1 DAL4A | 156.92 ± 4.65<sup>a</sup> | 121.83 ± 5.57<sup>b</sup> | 199.22 ± 8.51<sup>c</sup> | 153.42 ± 4.26<sup>a</sup> |
| 2 DAL4A | 203.75 ± 5.11<sup>a</sup> | 166.67 ± 5.35<sup>b</sup> | 253.89 ± 10.22<sup>c</sup> | 203.92 ± 5.75<sup>a</sup> |
| 3 DAL4A | 255.00 ± 5.41<sup>a</sup> | 204.83 ± 6.16<sup>b</sup> | 304.00 ± 8.72<sup>c</sup> | 259.25 ± 6.16<sup>a</sup> |
| 4 DAL4A | 309.00 ± 5.50<sup>a</sup> | 250.42 ± 6.67<sup>b</sup> | 349.22 ± 8.89<sup>c</sup> | 311.25 ± 6.25<sup>ac</sup> |
| 5 DAL4A | 362.92 ± 5.72<sup>a</sup> | 292.50 ± 7.26<sup>b</sup> | 400.56 ± 10.52<sup>c</sup> | 370.75 ± 7.19<sup>a</sup> |
| 6 DAL4A | 424.92 ± 5.91<sup>a</sup> | 319.42 ± 8.08<sup>b</sup> | 456.33 ± 11.00<sup>a</sup> | 430.58 ± 7.71<sup>a</sup> |
| 7 DAL4A | 478.33 ± 5.58<sup>a</sup> | 330.83 ± 8.02<sup>b</sup> | 510.11 ± 11.93<sup>a</sup> | 493.42 ± 7.59<sup>a</sup> |
| 8 DAL4A | 507.42 ± 5.59<sup>a</sup> | 337.33 ± 8.44<sup>b</sup> | 561.56 ± 11.61<sup>c</sup> | 543.92 ± 6.38<sup>c</sup> |
| 9 DAL4A | 521.25 ± 5.07<sup>a</sup> | 339.25 ± 8.40<sup>b</sup> | 613.11 ± 11.18<sup>c</sup> | 577.92 ± 6.70<sup>c</sup> |
| 10 DAL4A | 529.75 ± 5.18<sup>a</sup> | 339.92 ± 8.35<sup>b</sup> | 648.89 ± 10.22<sup>c</sup> | 601.25 ± 6.74<sup>d</sup> |
| 11 DAL4A | 535.08 ± 5.32<sup>a</sup> | 340.33 ± 8.29<sup>b</sup> | 669.00 ± 8.99<sup>c</sup> | 613.58 ± 6.24<sup>d</sup> |
| 12 DAL4A | 539.42 ± 5.38<sup>a</sup> | 340.33 ± 8.29<sup>b</sup> | 680.11 ± 9.95<sup>c</sup> | 619.58 ± 6.76<sup>d</sup> |
| 13 DAL4A | 541.00 ± 5.87<sup>a</sup> | 340.33 ± 8.29<sup>b</sup> | 686.89 ± 8.94<sup>c</sup> | 623.67 ± 7.44<sup>d</sup> |
| 14 DAL4A | 541.33 ± 5.92<sup>a</sup> | 340.33 ± 8.29<sup>b</sup> | 692.67 ± 8.84<sup>c</sup> | 626.42 ± 7.53<sup>d</sup> |
| 15 DAL4A | 541.58 ± 5.97<sup>a</sup> | 340.33 ± 8.29<sup>b</sup> | 696.00 ± 9.03<sup>c</sup> | 627.42 ± 7.65<sup>d</sup> |
| 16 DAL4A | 541.58 ± 5.97<sup>a</sup> | 340.33 ± 8.29<sup>b</sup> | 698.89 ± 9.21<sup>c</sup> | 627.67 ± 7.67<sup>d</sup> |

**Supplemental table T6:** Kinematic analysis of leaf 4 for a segregating *EF1ɑ:GIF1-07* population two days after leaf 4 appearance. . Significance was calculated by a two-tailed t test (n = 3). SEM = Standard error of the mean.

|  | Wild type | *EF1αGIF1-07* |  |  |  |
| --- | --- | --- | --- | --- | --- |
|  | Mean ± SEM | Mean ± SEM | P-value |  | % |
| Final leaf length (mm) | 553.61 ± 12.19 | 621.66 ± 6.97 | ✓ 1.33E-04 | ✓ | 10.94% |
| Leaf elongation rate (mm*h-1) | 2.12 ± 0.06 | 2.19 ± 0.09 | ✓ 0.52 | ✓ | 3.52% |
| Leaf elongation duration (h) | 237.69 ± 2.05 | 258.44 ± 2.55 | ✓ 7.64E-06 | ✓ | 8.02% |
| Division zone size (cm) | 1.13 ± 0.03 | 1.29 ± 0.01 | ✓ 2.66E-02 | ✓ | 12.59% |
| Cell division rate (cells cell-1 h-1) | 0.025 ± 0.002 | 0.023 ± 0.001 | ✓ 0.48 | ✓ | -10.33% |
| Cell cycle duration (h) | 27.31 ± 2.64 | 29.83 ± 2.06 | ✓ 0.49 | ✓ | 8.43% |
| Cell production (cell*h-1) | 15.77 ± 1.36 | 16.16 ± 0.90 | ✓ 0.82 | ✓ | 2.44% |
| Average size of dividing cells (μm) | 19.47 ± 1.51 | 20.04 ± 0.19 | ✓ 0.74 | ✓ | 2.81% |
| Mature cell size (μm) | 136.71 ± 11.91 | 137.07 ± 7.75 | ✓ 0.98 | ✓ | 0.26% |

**Supplemental table T7:**
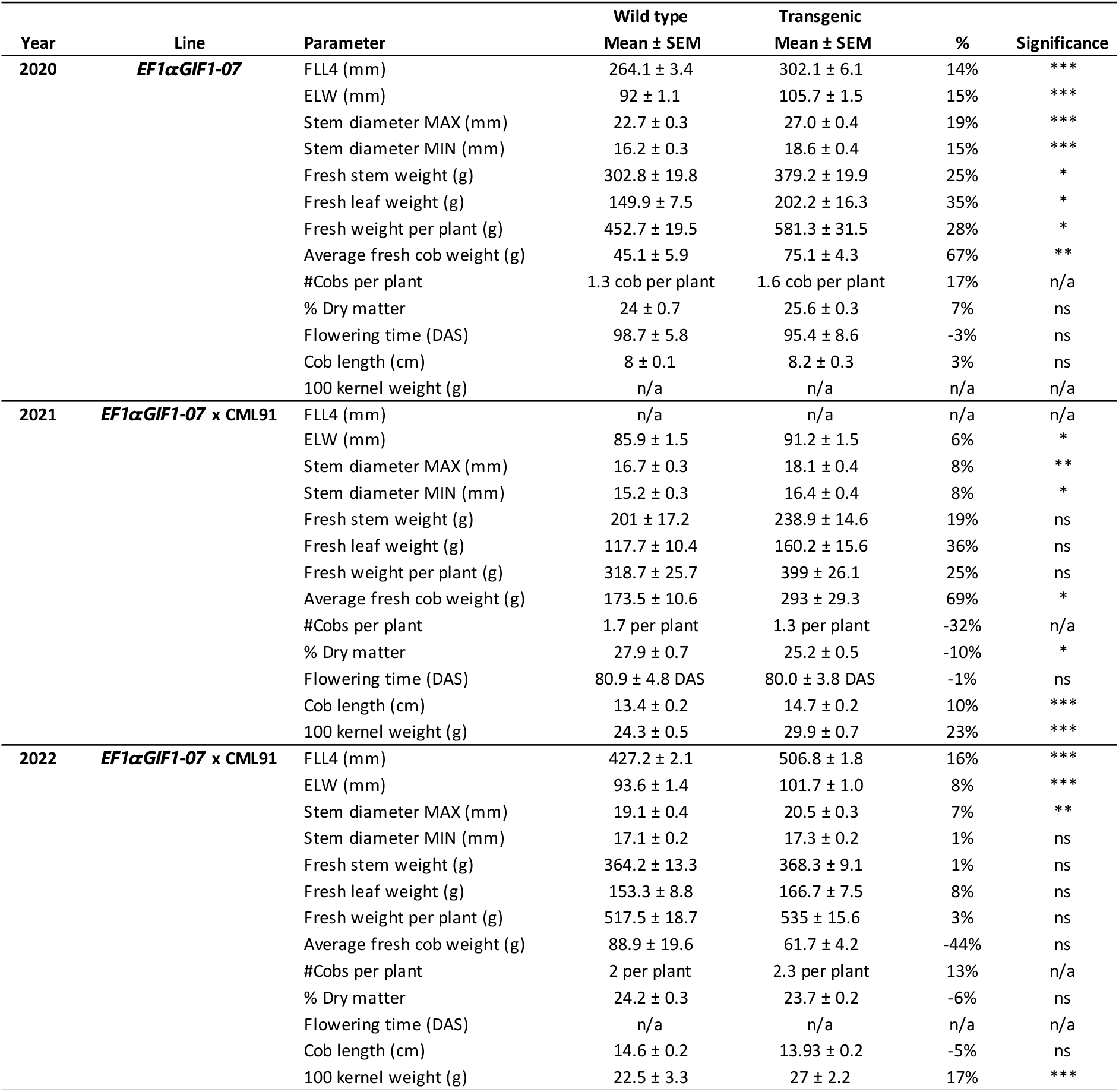
Growth, biomass and yield parameters in field trials of the *EF1ɑ:GIF1-07* and the *EF1ɑ:GIF1-07 x CML91* line. Significance was calculated by a two-tailed t test: * = P ≤ 0.05, ** = P ≤ 0.01, *** = P ≤ 0.001, ns = not-significant). SEM = Standard error of the mean. DAS = days after sowing. Leaf and biomass measurements were taken on 20 representatives plant per plot (n = 60). Flowering time was calculated based on the entire field. Cob length was measured for all obtained cobs (n ≥ 19 for *EF1ɑ:GIF1-07*, n ≥ 119 for *EF1ɑ:GIF1-07 x CML91* (year 1) and n ≥ 39 for *EF1ɑ:GIF1-07 x CML91* (year 2)). Seed measurements were performed on 5 representative cobs per plot (n = 15).

**Supplemental table T8:** Final leaf lengths of the *pla1-1;gif1-2* double mutants and segregating single *pla1-1* and *gif1-2* mutants in the greenhouse. Significance for final leaf number and final plant height was calculated by an analysis of variance with Bonferroni correction (n ≥ 3). Significance for final leaf length (leaf 9 - leaf 16) was calculated by a two-way analysis of variance with Bonferroni correction (n ≥ 3).

| | He-He<br>Mean ± SEM | *pla1-1*<br>Mean ± SEM | *gif1-2*<br>Mean ± SEM | *pla1-1;gif1-2*<br>Mean ± SEM |
| --- | --- | --- | --- | --- |
| Final leaf number | 16.86 ± 0.26<sup>ac</sup> | 19.57 ± 0.20<sup>b</sup> | 15.71 ± 0.18<sup>a</sup> | 6.00 ± 1.03<sup>c</sup> |
| Final plant height (mm) | 2005.00 ± 32.51<sup>a</sup> | 1816.00 ± 68.21<sup>a</sup> | 1189.00 ± 50.21<sup>b</sup> | 758.20 ± 117.7<sup>c</sup> |
| Final leaf length 9 (mm) | 880.57 ± 11.01<sup>a</sup> | 763.33 ± 21.49<sup>ab</sup> | 655.00 ± 22.47<sup>b</sup> | 496.80 ± 67.46<sup>c</sup> |
| Final leaf length 10 (mm) | 861.29 ± 15.05<sup>a</sup> | 778.00 ± 17.54<sup>ab</sup> | 657.63 ± 18.63<sup>b</sup> | 513.67 ± 89.35<sup>c</sup> |
| Final leaf length 11 (mm) | 858.00 ± 12.96<sup>a</sup> | 769.67 ± 21.08<sup>ab</sup> | 661.25 ± 15.03<sup>bc</sup> | 538.80 ± 89.06<sup>c</sup> |
| Final leaf length 12 (mm) | 837.00 ± 14.16<sup>a</sup> | 793.00 ± 28.46<sup>a</sup> | 635.25 ± 13.40<sup>bc</sup> | 538.40 ± 87.64<sup>c</sup> |
| Final leaf length 13 (mm) | 797.14 ± 15.22<sup>a</sup> | 783.14 ± 26.65<sup>a</sup> | 629.88 ± 13.91<sup>bc</sup> | 547.00 ± 97.24<sup>c</sup> |
| Final leaf length 14 (mm) | 756.86 ± 14.10<sup>a</sup> | 766.86 ± 22.32<sup>a</sup> | 564.13 ± 13.77<sup>bc</sup> | 514.20 ± 114.71<sup>c</sup> |
| Final leaf length 15 (mm) | 693.86 ± 19.23<sup>a</sup> | 760.14 ± 21.35<sup>a</sup> | 433.63 ± 38.65<sup>bc</sup> | 462.00 ± 137.63<sup>c</sup> |
| Final leaf length 16 (mm) | 520.29 ± 43.97<sup>a</sup> | 752.67 ± 25.01<sup>b</sup> | 334.17 ± 25.09<sup>c</sup> | 433.25 ± 128.17<sup>ac</sup> |

**Supplemental table T9:**
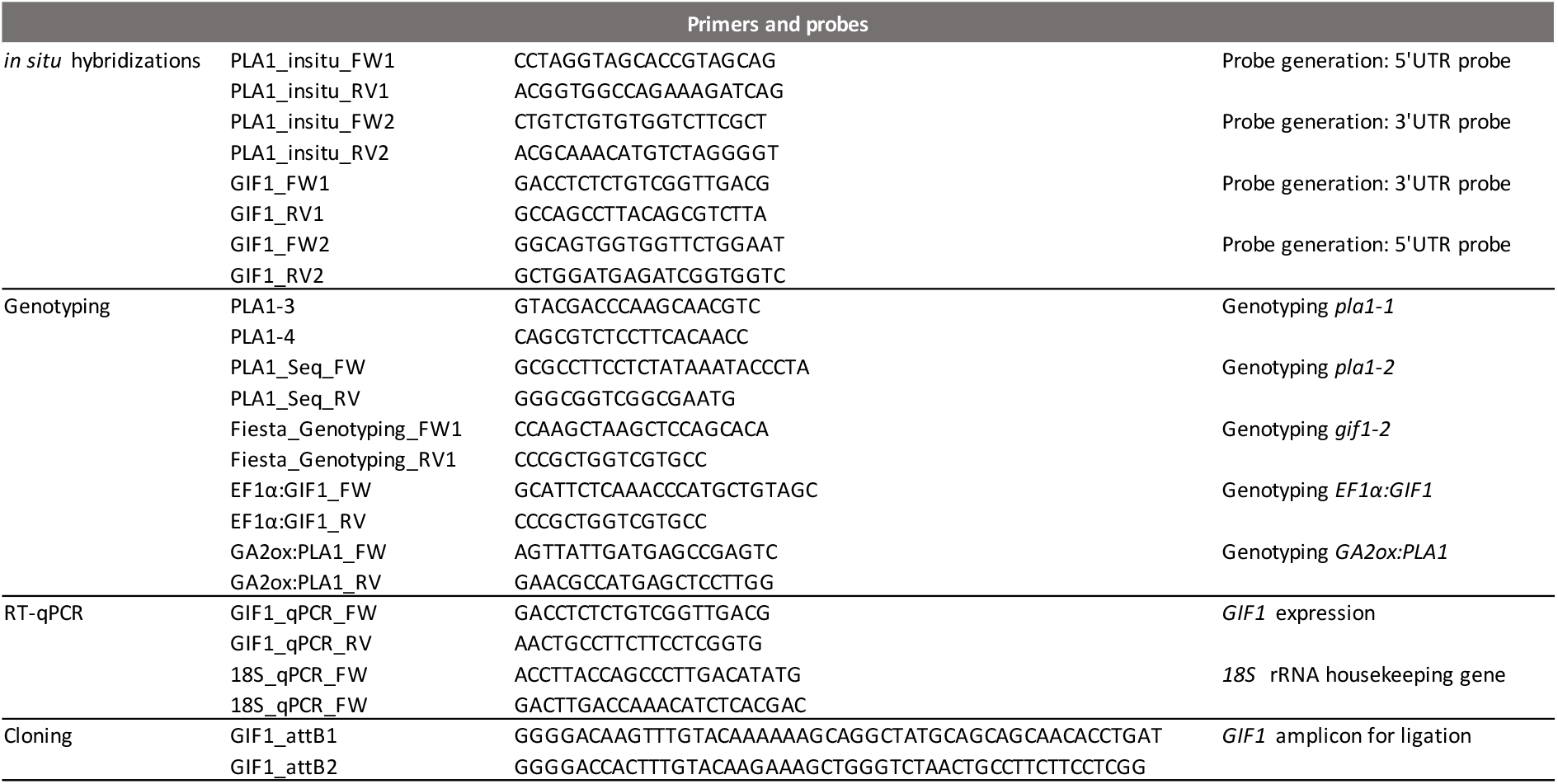
Primers and probes used in this study.

